# Sus1 maintains a normal lifespan through regulation of TREX-2 complex-mediated mRNA export

**DOI:** 10.1101/2022.04.07.487427

**Authors:** Suji Lim, Yan Liu, Byung-Ho Rhie, Chun Kim, Hong-Yeoul Ryu, Seong Hoon Ahn

## Abstract

Eukaryotic gene expression requires multiple cellular events, including transcription and RNA processing and transport. Sus1, a common subunit in both the Spt-Ada-Gcn5 acetyltransferase (SAGA) and transcription and export complex-2 (TREX-2) complexes, is a key factor in coupling transcription activation to mRNA nuclear export. Here we present that the SAGA DUB module and TREX-2 regulates distinctly yeast replicative lifespan in a Sir2-dependent and - independent manner, respectively. The growth and lifespan impaired by *SUS1* loss depend on TREX-2 but not on the SAGA DUB module. Notably, an increased dosage of the mRNA export factors Mex67 and Dbp5 rescues the growth defect, shortened lifespan, and nuclear accumulation of poly(A)^+^ RNA in *sus1Δ* cells, suggesting that boosting the mRNA export process restores the mRNA transport defect and damage in the growth and lifespan of *sus1Δ* cells. Moreover, Sus1 is required for the proper association of Mex67 and Dbp5 with the nuclear rim. Together, these data suggest that Sus1 links transcription and mRNA nuclear export to the lifespan control pathway, indicating that prevention of an abnormal accumulation of nuclear RNA is necessary for maintaining a normal lifespan.

## Introduction

Aging is a process accompanied by the gradual accumulation of molecular, cellular, and organ damage during sexual maturity, eventually leading to the decay of biological functions and increased vulnerability to morbidity and mortality [1, 2]. The budding yeast *Saccharomyces cerevisiae* is a useful model organism for studying the aging process, and this model is analyzed by with two distinct methods: replicative lifespan (RLS) and chronological lifespan (CLS) [3]. The RLS assay monitors how many daughter cells can be produced from a mother cell to study the lifespan of proliferating cells, such as stem cells, whereas CLS is a model of the aging process of postmitotic cells in multicellular organisms by measuring how long time a cell can survive in a nondividing state.

Lifespan studies using the yeast model system have revealed diverse conserved genetic pathways that influence aging, such as the Sir2 histone deacetylase-mediated maintenance of intact telomeric chromatin or suppression of rDNA recombination [4-7]. Among such factors involved in lifespan control, the Spt-Ada-Gcn5 acetyltransferase (SAGA) transcription coactivator complex, which has recently been identified as a regulator of the aging pathway, has multiple roles in the yeast lifespan. SAGA consists of four functionally independent modules: HAT module (histone acetylation), DUB module (deubiquitination of H2B), TAF module (coactivator architecture), and SPT module (assembly of the preinitiation complex) [8]. RLS is extended by the presence of a HAT inhibitor, inducing a similar effect as Sir2 activation, and is completely abolished by the loss of Gcn5, a subunit of the SAGA HAT module [9]. Furthermore, the heterozygous mutant *GCN5* or *NGG1*, a linker protein between Gcn5 and SAGA [10], leads to an increase in RLS [9]. However, yeast cells entirely lacking each component in the SAGA HAT module either do not exhibit an extended lifespan or rather show a decreased lifespan [11, 12]. Loss of Ubp8, Sgf73, or Sgf11 in the SAGA DUB module exceptionally extends RLS in a Sir2-dependent manner by enhancing telomeric silencing and promoting rDNA stability [11], whereas lacking the components in the SAGA SPT module mostly leads to a decrease in both RLS and CLS [12]. In particular, Spt7 in the SAGA SPT module is indispensable for a normal lifespan by maintaining genome stability and overall mRNA expression, and it is independent of Sir2 [12]. In addition, SAGA facilitates the retention and accumulation of extrachromosomal DNA circles and anchoring the DNA circle molecules to the nuclear pore complex (NPC), causing the organization of aged nuclei [13]. Although it is still uncertain how a single complex has multiple roles to ensure a normal lifespan, this is a good example of how aging is tuned finely by an intricate network of regulators.

SAGA interacts functionally and physically with the transcription and export complex-2 (TREX-2) complex composed of Sac3, Thp1, Cdc31, Sem1, and Sus1. This link is important for the transcription, mRNA export, and targeting of active genes to NPC [14]. The N-terminus in Sac3 provides a platform for association with Thp1 and Sem1 to form an mRNA-binding module and with the essential mRNA exporter Mex67, whereas its C-terminus binds to Sus1, Cdc31, and Nup1 nucleoporin, providing the attachment of the complex to NPC [15-19]. TREX-2 shares one subunit Sus1 with the SAGA DUB module, and Sus1 is also found at the promoter and coding regions in some SAGA-dependent genes, contributing to the coupling of transcription activation and mRNA export [20-23]. In particular, Sus1, Sac3, and Thp1 mediate the proper tethering of transcribed genes to NPCs upon the activation of transcription [24-26]. Both Cdc31 and Sem1 also contribute to promoting the association of TREX-2 with NPC and mRNA export [17, 27].

In eukaryotes, NPC, composed of ∼30 nucleoporin proteins, maintains a nuclear permeability barrier that selectively allows small molecules to diffuse in and out of the nucleus [28]. About half of the nucleoporins have characteristic domains with FG-repeat motifs, such as FG, FXFG, and GLFG. These FG domains are required for targeting nuclear membrane proteins, permeability barrier, and chromatin association with NPCs [29]. The age-dependent oxidative damage to nucleoporins disrupts the structure and function of NPCs in postmitotic cells, leading to the leakage of cytoplasmic proteins into the nucleus [30]. In entirely differentiated rat brain cells, nucleoporins are oxidized and long-lived without replacing newly synthesized proteins and their degradation, resulting in potentially harmful implications [30, 31]. Yeast RLS experiments provide more direct evidence of a correlation between NPCs and cellular lifespans [32]. Yeast cells lacking the GLFG domain of Nup116 exhibit a decrease in RLS. Such a shortened lifespan depends on Kap121-mediated defects in nuclear transport, which disrupt mitochondrial activity. In contrast, the Nup100-mediated nuclear export of specific tRNAs potentially limits yeast lifespan [32, 33].

Here we report that loss of *SUS1* triggers slow growth and a shortened lifespan in yeast cells. Although Sus1 certainly belongs to both the SAGA DUB module and TREX-2, cellular propagation and replicative ability impaired by *SUS1* loss are not altered by the additional deletion of the TREX-2 subunit, *sac3* or *thp1* but not the SAGA DUB module, suggesting that Sus1 is involved in the regulation of lifespan in a TREX-2-dependent manner. Also, unlike the SAGA DUB module, TREX-2-mediated lifespan control is independent of the presence of Sir2. Using a tiled yeast genomic DNA library, we found that overexpression of either Mex67 or Dbp5, mRNA export factors, suppresses growth defects in *sus1Δ* cells. Consistent with this result, decreased RLS and nuclear accumulation of poly(A)^+^ RNA upon the lack of *SUS1* were rescued by the increased dosage of Mex67 or Dbp5. Furthermore, Dbp5 mislocalization at the nuclear rim is greatly increased in *sus1Δ* cells. Taken together, these data suggest that Sus1 links transcription and mRNA nuclear export to the lifespan control pathway, indicating that blocking the abnormal accumulation of nuclear RNA is required for maintaining a normal lifespan.

## Results

### Deficient *sus1* allele results in the shortening of lifespan

Despite diverse attempts to define the correlation between the SAGA DUB module and yeast lifespan, the function of Sus1 in the lifespan remains still elusive [11]. To investigate the role of Sus1 in the control of yeast lifespan, we first analyzed the yeast RLS of the 50 cells from four independent *sus1Δ* strains, which exhibit comparable slow growth phenotype between strains (Figure 1A and B). Although McCormick and colleagues previously reported that loss of Sus1 had no effect on the lifespan when the replicative ability of 40 yeast cells was once examined by micromanipulation [11], we observed that the mean lifespans of all four independent *sus1Δ* strains were equally shorter than that of wild-type (WT) to a similar degree. The different result between the two studies is likely derived from the increased number of cells and bioreplicates in our study. These observations provided the first evidence that Sus1 is required for a normal cellular lifespan.

**Figure 1.**
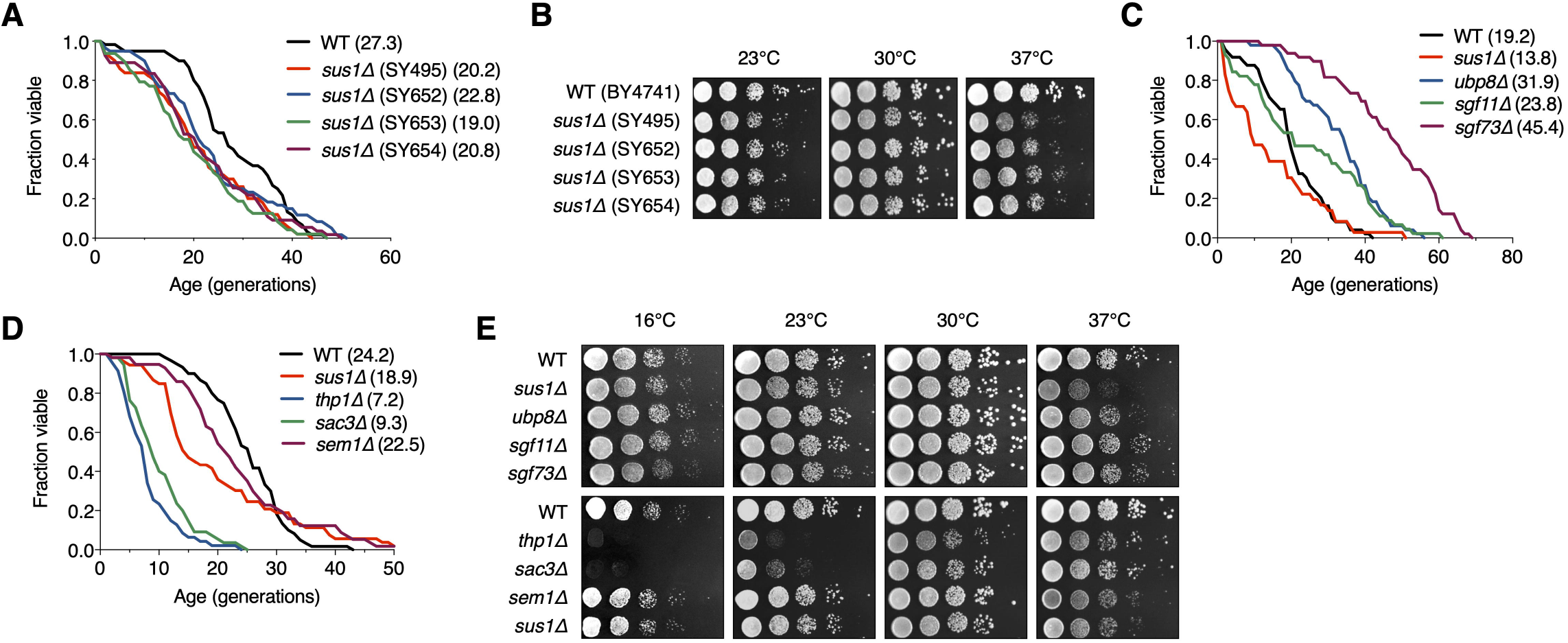
RLS is shortened by the loss of Sus1. (**A**) RLS analysis of four independent *sus1Δ* strains. The *SUS1* allele was replaced with *HISMX6* in SY495 and SY652 and *KanMX6* in SY653 and SY654, respectively. The mean lifespans are shown in parentheses. (**B**) Growth analysis of the strains used in (**C**). After spotting cells in 10-fold serial dilutions, the YPD plates were incubated at the indicated temperatures for 2–3 days. (**C** and **D**) RLS analysis of the indicated mutants of the SAGA DUB module (**C**) and TREX-2 (**D**). The mean lifespans are shown in parentheses. (**E**) Growth analysis of the strains used in (**C** and **D**). After spotting cells in 10-fold serial dilutions, the YPD plates were incubated for 2–3 days at 23°C, 30°C, and 37°C and for 5 days at 16°C.

### SAGA DUB module and TREX-2 affect differently lifespan

Sus1 is a component of the evolutionarily conserved SAGA DUB module and TREX-2 coupling of histone H2B deubiquitination-mediated transcriptional activation to nuclear pore-associated mRNA export [20, 21, 34]. Therefore, we next examined the change in RLS caused by the absence of nonessential genes encoding the individual subunits of the SAGA DUB module and TREX-2 (Figure 1C and D). Consistent with previous results [11], we observed exceptionally long-lived *ubp8Δ* and *sgf73Δ* strains and only a mild increase in the lifespan of the *sgf11Δ* strain, whereas the lack of these genes did not affect yeast cell growth (Figure 1C and E). In contrast, interestingly, loss of the two major structural components of TREX-2, Thp1 and Sac3, but not Sem1, impaired normal lifespan and vegetative growth at different temperatures more strongly than the lack of a linker protein Sus1 between the SAGA DUB module and TREX-2 (Figure 1D and E). Taken together, these results indicated that the SAGA DUB module and TREX-2 distinctly affect the cellular growth and lifespan, and the effects of their common subunit Sus1 correspond with that of TREX-2 but not the SAGA DUB module.

### Sus1 affects lifespan independently of other subunits in the SAGA DUB module

SAGA and TREX-2 complexes are physically and functionally linked, whereas their impact on the lifespan is clearly distinguished (Figure 1C-E). Therefore, we then sought to determine which complex-dependent pathways are responsible for shortened lifespan by loss of Sus1 (Figure 2). Sus1 is necessary for the association of Ubp8 and Sgf11 with the main body of SAGA, resulting in balanced H2B ubiquitination levels [21]. If a decreased lifespan in *sus1Δ* is due to the effects exclusively of the SAGA DUB module, Sus1 loss will not shorten the lifespan in *ubp8Δ, sgf11Δ*, and *sgf73Δ* backgrounds. However, the extended RLS of *ubp8Δ, sgf11Δ*, and *sgf73Δ* strains was significantly alleviated by the additional deletion of *SUS1*, serving Sus1 influences RLS through a distinct pathway with other components in the SAGA DUB module (Figure 2A-C). Furthermore, the slow growth phenotype and the heat and cold sensitivities of the *sus1Δ* strain were not affected by the additional deletion of *UBP8* or *SGF73*, although there is a mild increase of cold sensitivity in the *sus1Δ sgf11Δ* double mutant (Figure 2G). Overall, these data suggested that Sus1 contributes to the control of lifespan independently of the SAGA DUB module-mediated pathway.

**Figure 2.**
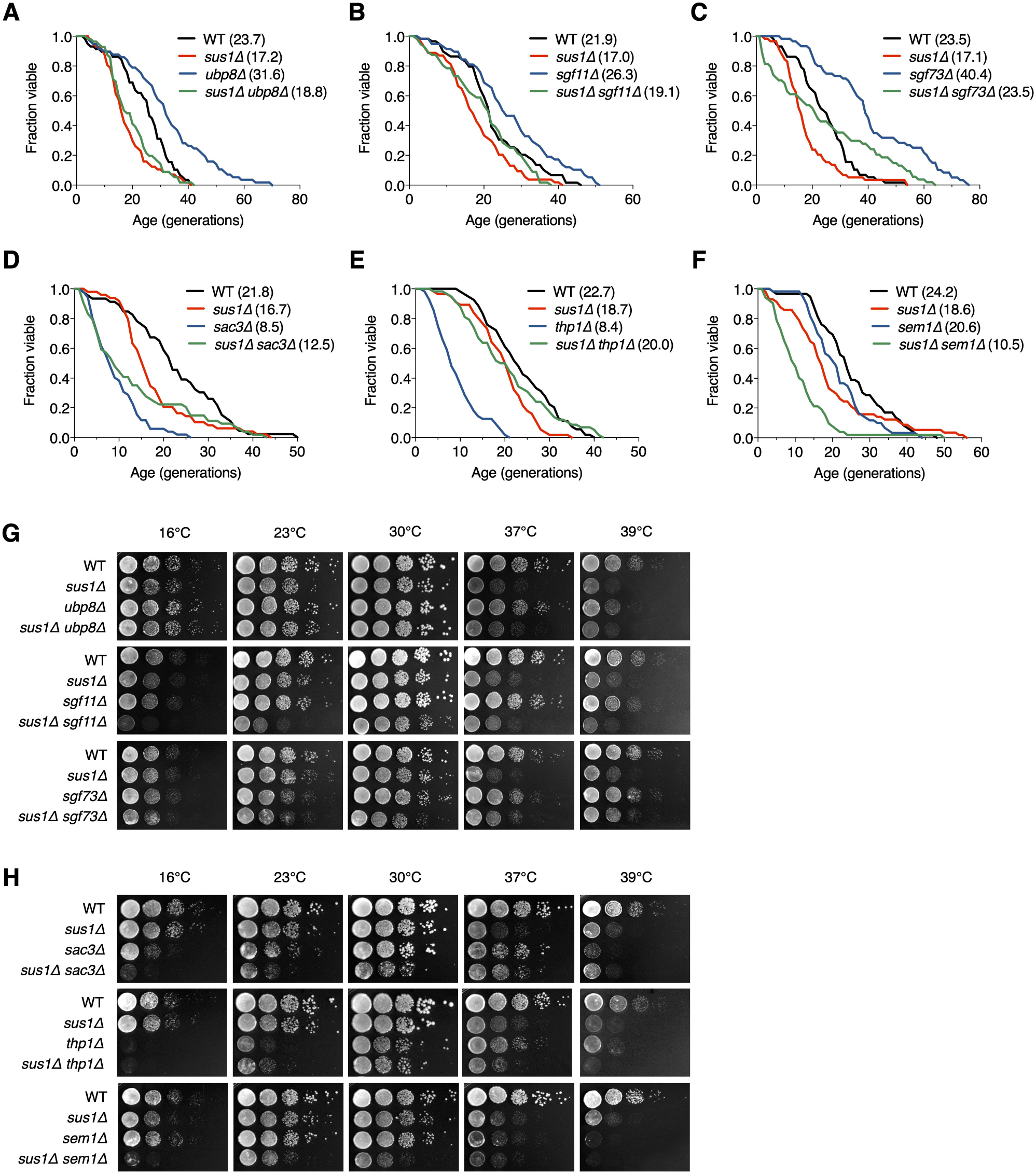
Additional mutations of TREX-2 components enhance RLS and growth defects of *sus1Δ*. (**A-F**) RLS analysis of the indicated mutants. The mean lifespans are shown in parentheses. (**G** and **H**) Growth analysis of the strains used in (**A-F**), as described in Figure 1E.

### Sus1 regulates lifespan through the TREX-2-dependent pathway

We next investigated whether Sus1-dependent lifespan shortening is determined by a TREX-2-driven pathway. Interestingly, additional *SUS1* loss did not affect the shortened lifespan in *sac3Δ*, but not *thp1Δ*, indicating that the effects of Sus1 on RLS relied on the presence of Sac3, which functions as a scaffold of TREX-2 (Figure 2D and E). RLS in the *sem1* mutant was indistinguishable from that in WT cells (Figure 1D). However, the altered lifespan of the *sus1Δ sem1Δ* double mutant significantly exceeded that of either single mutant, suggesting an additive effect on longevity as a result of combining deletion of *SUS1* with *SEM1* (Figure 2F). Although Sus1 and Sem1 are not the main and essential subunits of TREX-2, they effectively support TREX-2-mediated mRNP biogenesis and mRNA export [27, 34, 35]. Therefore, this result implied that the loss of both genes, *SUS1* and *SEM1*, boosts the suppression of TREX-2 activity. In contrast to the growth analysis results in *sus1Δ ubp8Δ* and *sus1Δ sgf11Δ* (Figure 2G), the slow growth and the heat, cold, and hydroxyurea (HU) sensitivities of *sus1Δ* were more significantly impaired by the additional loss of genes encoding the TREX-2 components, *SAC3, THP1*, or *SEM1*, suggesting that Sus1 is an important factor for TREX-2 activity (Figure 2H and supplementary Figure 1). Taken together, these results strongly suggested that Sus1 is necessary for normal cellular lifespan through a pathway dependent on TREX-2 but not the SAGA DUB module.

### TREX-2 functions in a Sir2-independent pathway

We next addressed how TREX-2 is involved in lifespan regulation. Because Sir2 is a well-known factor that modulates RLS [4, 6, 36], we subsequently investigated whether Sir2 affects RLS in TREX-2 mutants. It was previously reported that the SAGA DUB module, including Sus1, controls RLS via interaction with Sir2 [11]. As expected, we also observed that the extended lifespan in *ubp8Δ* and *sgf73Δ* failed due to the deletion of another gene (*SIR2*; Figure 3A). In addition, a similar result was found in *sir2Δ sgf11Δ*, which also failed to increase the lifespan by lacking Sgf11. However, although the decreased level of RLS was similar between those in *sir2Δ* and *sir2Δ sus1Δ* cells (Figure 3B), the absence of Thp1 or Sac3 more significantly impaired the shortened lifespan of *sir2Δ* cells (Figure 3C), suggesting that TREX-2 has a role in lifespan modulation via a Sir2-independent mechanism. Because *sus1* deletion has a milder effect on RLS than *thp1* or *sac3* mutations (Figure 1D), Sus1 might marginally affect a change in the lifespan of the *sir2Δ* strain.

**Figure 3.**
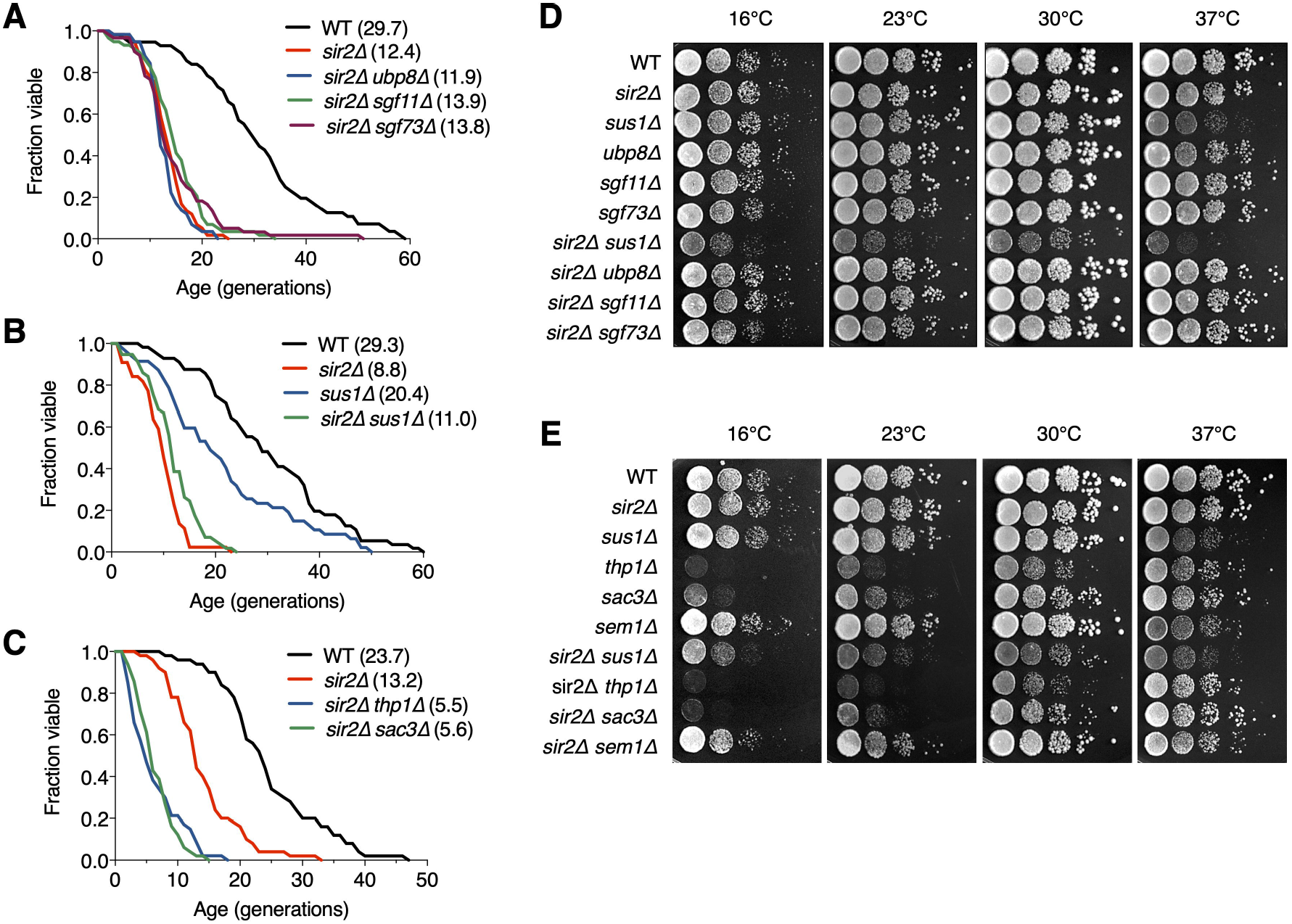
TREX-2 affects RLS in a Sir2-independent manner. (**A-C**) RLS analysis of double deletion strains of SAGA DUB mutants (**A**), *sus1Δ* (**B**) and TREX-2 mutants (**C**) in combination with *sir2Δ*. The mean lifespans are shown in parentheses. (**D** and **E**) Growth analysis of the strains used in (**A-C**), as described in Figure 1E.

The silencing function of Sir2 contributes to extending RLS by suppressing rDNA recombination [4]. Nevertheless, contrary to the lack of Ubp8, loss of Sus1 affected neither silencing nor recombination at the rDNA loci (supplementary Figure 2). Instead, *SIR2* loss significantly impaired the growth of *sus1Δ* cells, whereas Sir2 did not affect the growth of mutants of other components in the SAGA DUB module or TREX-2 (Figure 3D and E), suggesting that at least Sus1 influences yeast cell growth in a Sir2-independent pathway.

Taken together, these data indicated that the SAGA DUB module and TREX-2 are required for lifespan regulation through distinct cellular mechanisms. In particular, this reveals a Sir2-independent novel pathway to sustain RLS, perhaps by TREX-2 activity.

### Overexpression of the nuclear RNA export factors Mex67 and Dbp5 rescues the lifespan defect in *sus1Δ*

TREX-2 associates with not only SAGA but also with the NPC basket structure, and its ability to interact with multiple factors contributes to establishing the link between transcription, pre-mRNA processing, and mRNA export [37]. Age-dependent deterioration of NPCs leads to loss of the nuclear permeability barrier, causing defects in nuclear integrity [30]. Each FG nucleoporin, an intrinsically disordered regulator of nucleocytoplasmic transport, in NPC distinctly affects aging, whereas Nup100 loss shows increased lifespan, and longevity is impaired in *nup16* mutants [32, 33].

To determine whether the TREX-2-mediated lifespan regulation pathway is related to the role of NPC in the aging control mechanism, we screened for NPC genes that prevent a defect in cell growth, which is usually positively related to RLS, upon *SUS1* deletion (Figure 4B and supplementary Figure 3). To monitor cell proliferation immediately after a *SUS1* gene was lost, we used a strain carrying the *sus1Δ* allele in the chromosome and a covering WT *SUS1* plasmid that could be readily evicted on 5-FOA medium; the *URA3* gene product is toxic to cells on 5-FOA (Figure 4A). The WT strain (*sus1Δ* + p*SUS1*) was then transformed with 46 different library plasmids containing individual genes of NPC and proteins known to physically interact with NPC, searched in the *Saccharomyces* Genome Database (SGD; https://www.yeastgenome.org), and yeast cell growth was measured after evicting the *SUS1* cover plasmid (Figure 4B and supplementary Figure 3). Transformants expressing a library plasmid, including Asm4, an FG nucleoporin component of the central core of NPC [38], displayed a more severe growth defect than that of the null *sus1* mutant, implying a cooperation between TREX-2 and the previously reported role of NPC in aging. More interestingly, additional copies of plasmids expressing the mRNA export machinery Mex67 or Dbp5 (Figure 4C) significantly suppressed growth reduction in *sus1Δ*, although there were no effects of plasmids, including *MTR2* or *NAB2*, which are genes encoding a cofactor of Mex67 or a physiological target of Dbp5, respectively [39, 40], on the growth of the *sus1Δ* strain. Furthermore, it was previously reported that *sus1Δ* is synthetically lethal with *dbp5* or *mex67* mutant alleles [20]. The *MEX67*-DAmP strain, in which mRNA destabilization of an essential gene is generated by disruption of its natural 3′-untranslated region [41], but not *MTR2*-DAmP, showed a shortened lifespan (Figure 4D). Strikingly, consistent with the growth assay results in Figure 4B, the existence of library plasmids encoding *MEX67* or *DBP5* significantly restored the decreased lifespan of the *sus1Δ* strain (Figure 4E). Moreover, the strong overexpression of the single *MEX67* gene rescued cell proliferation and RLS impaired by *SUS1* loss (Figure 4F and G). In contrast to the growth and RLS results obtained from a library plasmid expressing *DBP5* (Figure 4B and E), Dbp5 expressed from pRS425 only improved the shortened lifespan, but not the impaired growth rate, in *sus1Δ* (Figure 5F and G). Presumably, a proper dosage of *DBP5* is required to entirely overcome the cellular stress caused by Sus1 loss. Taken together, these results suggested that an increased dosage of the mRNA export factors Mex67 and Dbp5 helps circumvent the growth and lifespan defects derived from Sus1 loss.

**Figure 4.**
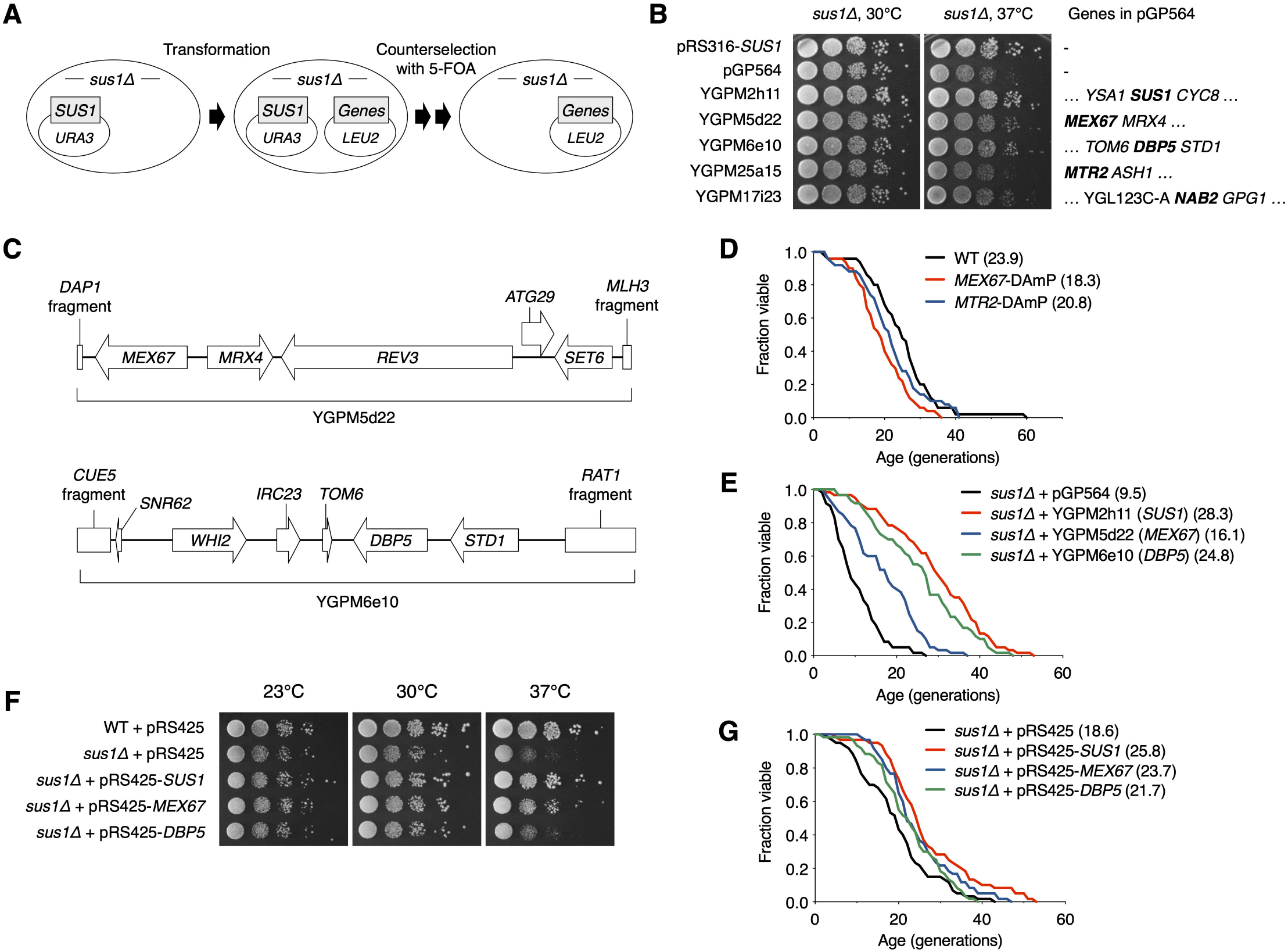
Multicopies of *MEX67* or *DBP5* rescue impaired RLS in *sus1Δ*. (**A**) Strategy used to identify specific genes that suppress *sus1Δ* defects. *sus1Δ* cells containing pRS316-SUS1 were transformed with the indicated pGP564 (*LEU2*)-based plasmids, including NPC-related genes. Cells were streaked on SC-Trp-His-Leu supplemented with 5-FOA twice to evict pRS316-SUS1. (**B** and **F**) Growth analysis of WT or *sus1Δ* strains including the indicated plasmids, as described in Figure 1E. Each gene on the plasmids is listed on the right panel of (**B**). (**C**) Schematic diagrams of plasmids YGPM5d22 (top) and YGPM6e10 (bottom). The ORF locations (arrows or boxes) and gene names are indicated. (**D, E** and **G**) RLS analysis of the indicated mutants (**D**) and *sus1Δ* cells, including the indicated plasmids (**E** and **G**). RLS analysis in (**E**) or in (**G**) was carried out on SC-leu or SC plates, respectively. The mean lifespans are shown in parentheses.

**Figure 5.**
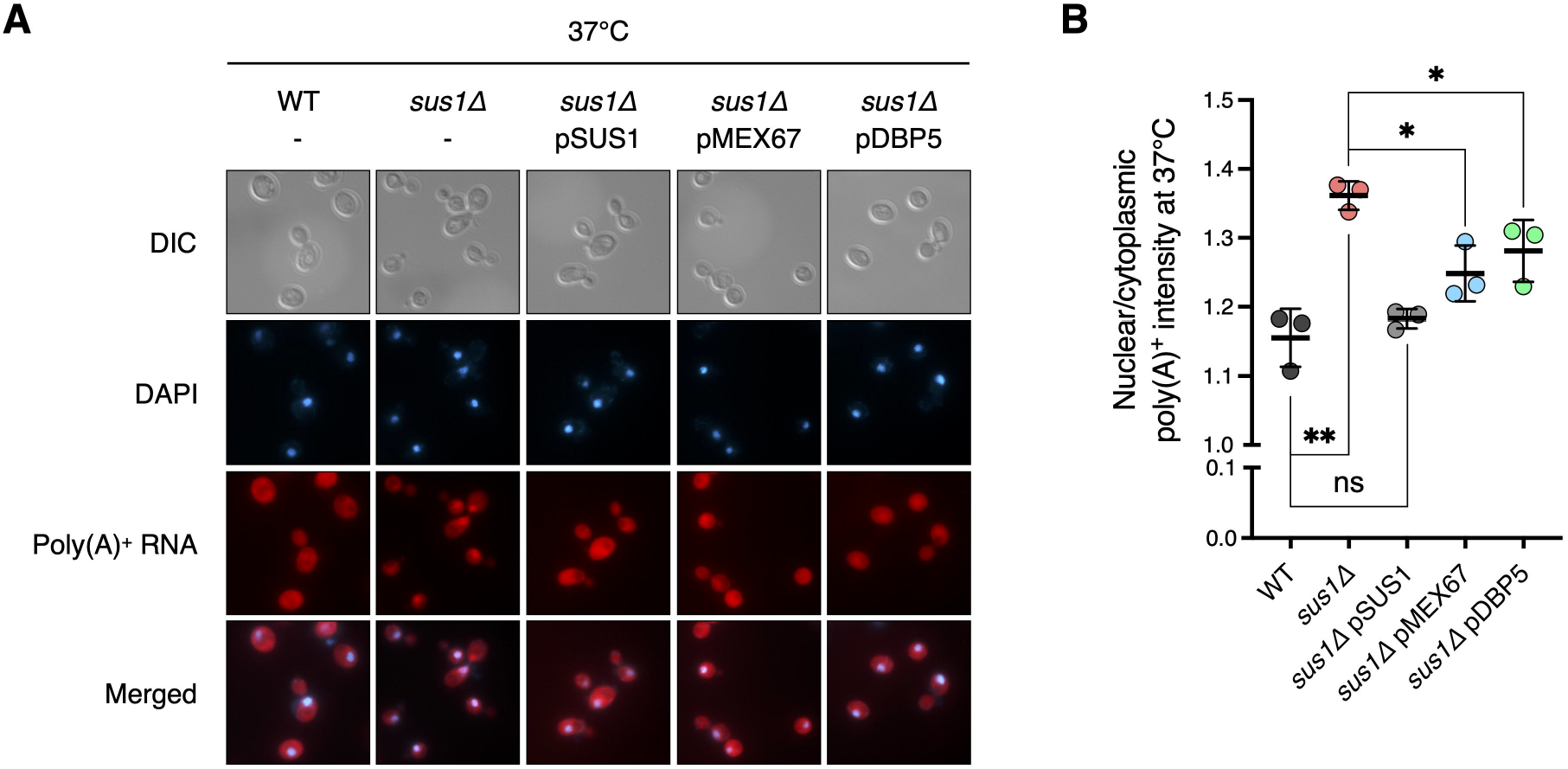
Increased doses of *MEX67* or *DBP5* restore the mRNA export defect in *sus1Δ*. (**A**) Representative images of FISH analysis for the WT and *sus1Δ* containing pRS316 expressing the indicated genes. Poly(A)^+^ RNAs were hybridized with Cy3-labeled oligo(dT) probes and visualized by fluorescence microscopy. DNA was stained by DAPI. DIC, DAPI, poly(A)^+^ RNA, and merged images are shown. (**B**) Quantitative analysis of FISH results in (**A**). The nuclear poly(A)^+^ RNA intensity of each cell was divided by the cytoplasmic poly(A)^+^ RNA signal of the corresponding cell. Data are the mean ± SD of triplicate experiments. **, *P* < 0.01; *, *P* < 0.05; ns, not significant (Student’s *t*-test between the indicated pairs of values).

### Increased *MEX67* or *DBP5* expression rescues the mRNA export defect in *sus1Δ*

Mutants of various factors involved in mRNA export, such as Sus1, induce nuclear retention of mRNA transcripts with poly(A)^+^ tails and sequester newly synthesized transcripts within nuclear foci at or near transcription sites [20, 42]. Given that multiple copies of the mRNA export factors Mex67 and Dbp5 compensate for the lifespan defect in *sus1Δ*, we next carried out poly(A)^+^ RNA fluorescence *in situ* hybridization (FISH) to determine whether Mex67 or Dbp5 would rescue the mRNA export defect of *sus1Δ* (Figure 5). Consistent with previous results in *sus1Δ* cells [20], we observed a nuclear accumulation of poly(A)^+^ RNA in cells lacking *SUS1* at 30°C, and the signal of the nuclear poly(A)^+^ signal became even stronger by transferring cells to 37°C, probably reflecting the overproduction of heat shock mRNAs and an even more severe mRNA export defect (supplementary Figure 4). In contrast to the impaired mRNA export in *sus1Δ* cells with an empty vector, mRNA export was fully rescued in WT *SUS1* gene-reintroduced *sus1Δ* cells (Figure 5A, lanes 1–3 and supplementary Figure 5). Significantly, an extra copy of Mex67 or Dbp5 suppresses the accumulation of poly(A)^+^ RNA at nuclear foci in *sus1Δ* (Figure 5A, lanes 4 and 5 and supplementary Figure 5), supporting the hypothesis that the enhanced mRNA export by an increased dosage of Mex67 or Dbp5 restores the mRNA export defect and impaired lifespan in cells lacking *SUS1*.

### Sus1 is required for Dbp5 localization at the nuclear rim

Finally, we determined why the *sus1* deletion-mediated inhibition of mRNA nuclear export was reversed by the additional presence of general mRNA export factors. Mex67 and its heterodimeric partner Mtr2 are generally localized to the entire nuclear rim [43]. However, the subcellular location of Mex67 was altered and focused exclusively on some sites at the nuclear rim in *sac3* mutants [34, 44]. Also, Mex67 localization was partially affected by Sus1 loss [34]. Therefore, to evaluate the effects of Sus1 on positioning mRNA export factors at the nuclear rim, we monitored the nuclear location of green fluorescent protein (GFP)-tagged Dbp5 by fluorescence microscopic analysis (Figure 6). As expected, strong foci generated by Dbp5-GFP mislocalization at the nuclear rim increased greatly in *sac3Δ* cells. In particular, the ratio of GFP intensity of strong focus to the rest of the nuclear rim in *sus1Δ* was elevated to a level similar to that in *sac3Δ*. However, *sus1Δ sac3Δ* double mutants did not show any additional defect, suggesting that the two genes are epistatic and that mRNA export factor mislocalization in *sus1Δ* may be a critical reason for its accumulation of nuclear poly(A)^+^ RNA. Therefore, these results, together with RLS (Figure 4) and poly(A)^+^ RNA FISH (Figure 5) data, strongly suggested that Sus1 is required for maintaining a normal lifespan through mRNA export control.

**Figure 6.**
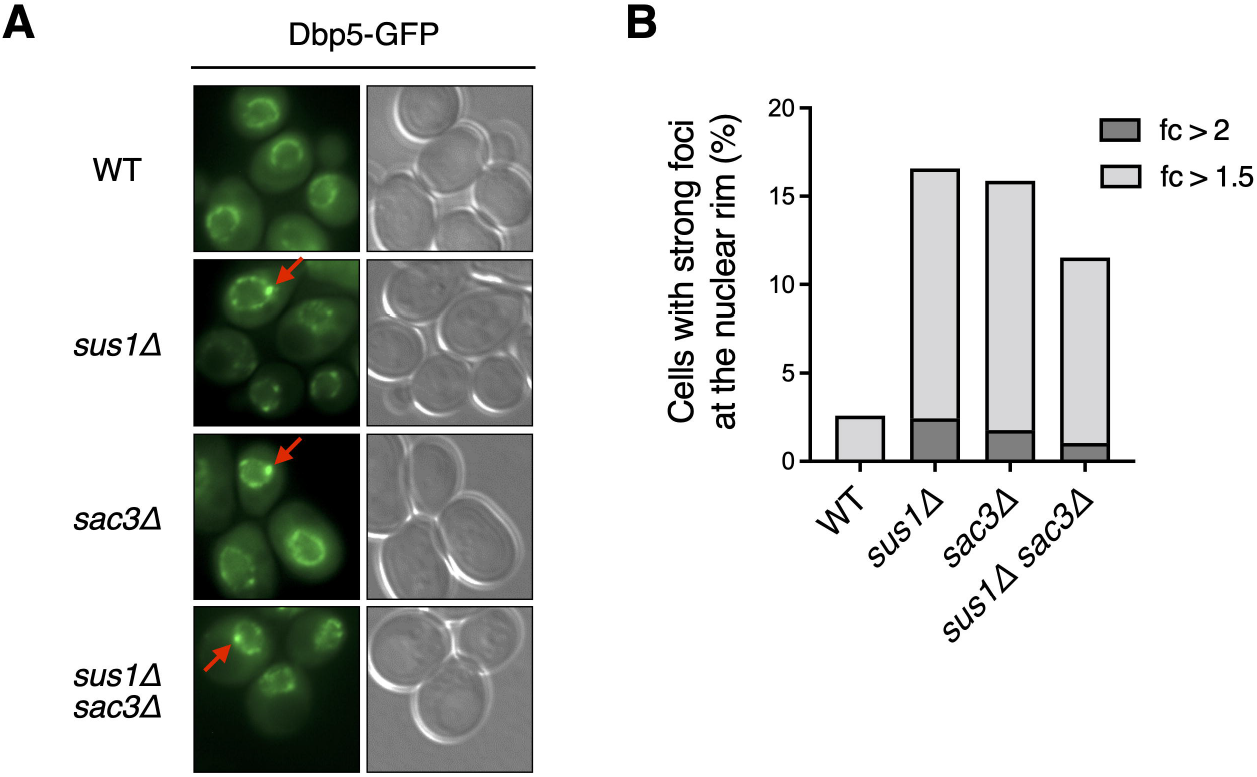
Dbp5 is mislocalized in *sus1Δ* cells. (**A**) Fluorescence microscopic analysis of Dbp5-GFP in WT and the indicated mutants. The left and right panels show the GFP and DIC images, respectively. Red arrows indicate strong foci at the nuclear rim. (**B**) Percentage of cells containing strong foci of Dbp5-GFP shown in (**A**). The ratio of GFP intensity of strong focus to that in the outer region of strong focus is calculated. Light and dark gray color bars in the graph indicate the percentage of cells with >1.5- and 2-fold of the focus ratio, respectively.

## Discussion

We revealed that Sus1, a cosubunit of SAGA and TREX-2, is a novel regulatory factor in the lifespan regulation in yeast. Although many factors affect yeast lifespan in a Sir2-dependent manner, Sus1-mediated maintenance of the normal lifespan is dependent on NPC-associated TREX-2 but not on SAGA. An mRNA export defect caused by *SUS1* loss is restored by an increased dosage of the NPC anchoring-dependent mRNA export factors Mex67 and Dbp5, resulting in the rescue of RLS in *sus1Δ*. Furthermore, Sus1 is necessary for the cellular localization of Mex67 and Dbp5. Therefore, Sus1 integrates two processes, mRNA export and lifespan control, to maintain the youthful state of the nucleus by blocking the abnormal accumulation of mature mRNA in the nucleus.

SAGA plays multiple roles in transcription, including histone modification and RNA metabolism [14], and the individual loss of its components differently affects RLS in yeast [9, 11, 12], implying that the complex network of subunits needs to ensure a normal lifespan. Additionally, RLS is distinctly influenced by null mutants of nucleoporin proteins [32, 33]. NPC is composed of diverse proteins that form the transport barrier [28], and its deterioration is considered as an aging marker [30, 31]. The cellular lifespan is a complex concept governed by multiple factors, and understanding such complex interactions is crucial for developing a new therapeutic strategy for age-related pathologies [45, 46].

Transcription accelerates the DNA damage rate, which can cause symptoms of genome instability and further premature aging [47]. An error rate of transcription increases with aging and induces the aggregation of peptides associated with age-related disorders [48]. Additionally, diverse transcription factors can act as regulators of lifespan in eukaryotes, reflecting a link between gene expression and aging [49-57]. Notably, nuclear events after transcription are certainly related to cellular aging. In *Drosophila*, sensitivity to environmental stress is increased, and lifespan is reduced by the mutation of the THO complex involved in transcription elongation and mRNA export [58]. The decreased turnover rate of human RNA by oxidative stress or reduced RNA exosome activity is one of the main causes of cellular senescence [59]. *PHO84* gene expression is repressed by its antisense RNA transcript in chronologically aged yeast cells, and antisense RNA stabilization is governed by Rrp6 exosome and histone acetylation [60]. To prevent pathophysiological cell senescence and cell death, therefore, it is important to identify mechanisms that inhibit transcription errors, nuclear accumulation of RNA, and abnormal export of mature RNA. The newly identified aging factor Sus1 is conserved in higher eukaryotes and appears to be important for maintaining the optimal balance of such mechanisms.

## Materials and methods

### Yeast strains

The yeast strains used in this study are listed in supplementary Table 1. Standard techniques were used for strain construction. The deletion strains were generated by replacing each open reading frame (ORF) with *KanMX* modules constructed by polymerase chain reaction (PCR) amplification from the corresponding strains obtained from EUROSCARF or pFA6a-KanMX6 or *HIS3MX6* module derived from pFA6a-HIS3MX6 [61]. The SY1031 strain was generated by switching *HIS3MX6*, used as a marker of DBP5-GFP in FY740, to the *KlURA3* obtained from pFA6a-GFP-KlURA3 (Sung et al. 2008). To generate the SY1035 strain, the C-terminal insertion cassette of *SUS1-HA* was constructed by PCR amplification from pFA6a-HA-KlURA3 [62] and transformed into BY4741 cells. For the SY1036 and SY1037 strains, *MEX67-GFP* of FY739 and *DBP5-GFP* of FY740 were individually swapped by C-terminal HA tagging cassettes derived from pFA6a-HA-KlURA3 [62]. All strains were verified by PCR and/or immunoblot analysis.

### Plasmids

The plasmids used in this study were created as described previously [63] and are listed in supplementary Table 2. To make pRS316-SUS1, pRS316-MEX67, and pRS316-DBP5, ORFs of *SUS1, MEX67*, and *DBP5*, including 900 bp upstream and 900 bp downstream, were individually PCR-amplified from yeast genomic DNA (BY4741) and cloned into pRS316. To create pRS425-SUS1, pRS425-MEX67, and pRS425-DBP5, *SUS1, MEX67*, and *DBP5* tagged with a sequence for the triplicated HA epitope, including 900 bp upstream and 700 bp downstream of the ORF, was PCR-amplified from SY1035, SY1036, and SY1037, respectively, and cloned into pRS425. All constructs were confirmed by DNA sequencing.

### Spotting assay

The spotting assay was performed as described previously [64]. Liquid cultures in exponential growth were normalized to 0.1 OD_600_ and subjected to 10-fold serial dilutions. Cells were spotted onto YPD medium with or without 150 mM HU or SC medium with the appropriate amino acids and bases, and the plates were incubated at 16°C, 23°C, 30°C, or 37°C for 2–5 days. For *URA3*-based rDNA silencing analysis, strains containing the *URA3* gene at rDNA loci were used as described previously [65]. The pregrown cells were normalized to 1.0 OD_600_ and then diluted with 5-fold serial dilutions. Cells were spotted onto SC medium with or without uracil or containing 5-FOA, and the plates were incubated at 30°C for 2–3 days.

### RLS analysis

The RLS of yeast strains was measured on YPD plates, unless otherwise indicated, as described previously [12, 66]. A total of ∼50 virgin daughter cells were subjected to lifespan analysis. To assess lifespan differences, a Mann-Whitney test was performed with a cutoff of *P* = 0.05. The average lifespan was considered different when *P* < 0.05 [4]. The comparison values between the control and each mutant are listed in supplementary Table 3.

### Fluorescence microscopic analysis

The images were acquired using an AxioCam HRm mounted on an Axio Observer. Z1 microscope with a Plan-Apochromat 100×/1.40 Oil DIC M27 objective lens, as reported previously [12]. Zeiss Filter Sets 38 HE (489038-9901-000), 20 (488020-9901-000), and 49 (488049-9901-000) were used to observe the fluorescence of GFP (excitation, 488 nm; emission, 509 nm), Cy3 (excitation, 549 nm; emission, 562 nm), and 4′,6-diamidino-2-phenylindole (DAPI; excitation, 359 nm; emission, 463 nm), respectively. ZEN 2012 Blue Edition software was used to acquire and process each fluorescence image.

Dbp5-GFP foci formation was determined as described previously [34, 44]. The GFP fluorescence intensity was measured using ImageJ software. Each GFP intensity of strong focus was divided by that of the outer region of strong focus in the corresponding cell, and the calculated ratio was collected from at least 1,000 cells per strain.

### Poly(A)^+^ RNA FISH

Poly(A)^+^ RNA FISH was performed as described previously with minor modifications [67]. Yeast cells were grown in 10 ml SC-Ura medium at 30°C to 0.3 OD_600_. Each culture was then divided into two halves and incubated for 2 h at 30°C or 37°C, respectively. Cells were crosslinked by adding 1:10 volume of 37% (w/v) formaldehyde and incubated for 1 h at room temperature. Crosslinked cells were washed twice with 0.1 M potassium phosphate buffer (pH 6.5) and once with 1.2 M sorbitol/0.1 M potassium phosphate buffer (pH 6.5). The washed cells were allowed to adhere on a 0.3% poly-L-lysine (Sigma)-coated 10-well slide. The cell wall was digested by treating 250 μg/ml Zymolyase 20T (MP Bio) for 20 min. Cells were washed and hybridized with hybridization solution [50% deionized formamide, 4× SSC, 1× Denhardt’s solution, 125 μg/ml tRNA (Sigma), 10% dextran sulfate, 500 μg/ml denatured salmon sperm DNA (Sigma), 50 pM Cy3-labeled oligo(dT)_30_ (custom□synthesized by Integrated DNA Technologies)] overnight at 37°C in a humid chamber. After hybridization, cells were washed, air-dried, and mounted with mounting solution [70% glycerol, 1× PBS, 1 mg/ml p-phenylenediamine, and 1 μg/ml DAPI (Sigma)]. ImageJ was used to detect nuclear and cytoplasmic Cy3 signals. At least 100 cells were examined for each of the three independent experiments. The significance between the indicated strains was determined by a two-tailed, unpaired Student’s *t*-test using GraphPad Prism (**, *P* < 0.01; *, *P* < 0.05). Data represent the mean ± standard deviation (SD) of triplicate experiments.

### Unequal sister chromatid exchange assay

The rate of marker loss through the unequal recombination of an *ADE2* marker inserted into the rDNA array was measured as reported previously [12, 65]. Cells grown to 1.0 OD_600_ were plated at a density of ∼400 cells per SC plate containing a low adenine concentration (27 μM). The plates were incubated at 30°C for 2 or 3 days and then stored at 4°C for ∼2 weeks to enhance the red color development. Colonies were counted using GeneTools software (Syngene). The percentage of marker loss was calculated by dividing the number of red-sectored colonies by the total number of colonies. Completely red colonies, indicating marker loss before plating, were excluded from the calculation.

## Supporting information

Supplementary Figure S3 (continued)

## Authors’ Contributions

S.H.A. organized and designed the scope of the study. H.-Y.R. wrote the manuscript with assistance from S.L., C.K. and S.H.A. S.L., Y.L. and B.-H.R. performed spotting assay and/or lifespan analysis. S.L. and Y.L. created plasmids used in this study. S.L. performed spotting assays using a tiled yeast genomic DNA library, unequal sister chromatid exchange assays and all fluorescence microscopic analyses. S.L. and S.H.A. performed data analysis for all experiments.

## Acknowledgments

We thank W.-K. Huh, S.-T. Kim, and M.S. Longtine for supplying the yeast strains and DNA constructs.

