## Supplementary Materials for "Sus1 maintains a normal lifespan through regulation of TREX-2 complex-mediated mRNA export"

**Supplementary Figure legends**

**Supplementary Figure 1. HU sensitivity assays in double deletion strains of *sus1Δ* with TREX-2 mutants.** The indicated mutants were spotted onto YPD plates with or without 150 mM HU, as described in Figure 1E.

**Supplementary Figure 2. *SUS1* deletion does not affect rDNA silencing and recombination.** (**A**) The schematic diagram (top) shows an rDNA unit embedded within a tandem array on chromosome XII with the position of *mURA3* reporters inserted into NTS1 or NTS2. The 35S pre-rRNA encoding the 18S, 5.8S, and 25S rRNAs is separated by NTS1 and NTS2. The locations of RFB (double triangle), ARS replication origin (oval), 5S rRNA gene (triangle), and 35S transcription start site (bent arrow) are represented. *URA3*-based rDNA silencing assays (bottom) were carried out in DMY2798 (*leu2::mURA3*), DMY2804 (*RDN1-NTS1::mURA3*), or DMY2800 (*RDN1-NTS2::mURA3*) strains with the indicated deletions. (**B**) The frequency of unequal rDNA crossovers was monitored by loss of the *ADE2* gene located within the rDNA array for WT (W303R) and the indicated deletion strains. Pictures of plates are shown in the top panels. The percentage of *ADE2* gene loss (% marker loss) was calculated as the ratio of red-sectored colonies to the total number of colonies and shown in the bottom panel. Completely red colonies were excluded.

**Supplementary Figure 3. Screening of NPC-related genes for the suppression of growth defects in *sus1Δ*.** Growth analysis of *sus1Δ* strains, including the indicated plasmids, as described in Figure 1E. Genes on the plasmids (pGP564) are listed on the right of each panel.

**Supplementary Figure 4. The mRNA export defect by *sus1Δ* is observed at both 30℃ and 37℃.** (**A**) Poly(A)^+^ RNA FISH analysis of WT, *sus1Δ*, and *sus1Δ* containing pRS316-SUS1, as described in Fig. 5A. (**B**) Violin plot of poly(A)^+^ RNA FISH results of strains used in Figure 5A at both 30℃ and 37℃. The medians and quartiles are marked as thick and dotted lines, respectively. ****, *P* < 0.0001; **, *P* < 0.01; *, *P* < 0.05 (Student’s *t-*test between the indicated pairs of values).

**Supplementary Figure 5.** **Additional copies of *MEX67* or *DBP5* rescue the mRNA export defect in *sus1Δ*.** (**A**) Violin plot of poly(A)^+^ RNA FISH results of Figure 5A, as described in supplementary Figure 4B. The nuclear/cytoplasmic poly(A)^+^ intensity of each replicate is plotted. (**B**) The cell numbers and mean intensity of nuclear/cytoplasmic poly(A)^+^ in (**A**) are shown.

**Supplementary Table 1.** Strains used in this study

| Strain | Genotype | Source |
| --- | --- | --- |
| BY4741 | *MATa ura3∆0 leu2∆0 his3∆1 met15∆0* | EUROSCARF |
| FY231 | *MATa ura3∆0 leu2∆0 his3∆1 met15∆0 ubp8∆::KanMX4* | EUROSCARF |
| FY390 | *MATa ura3∆0 leu2∆0 his3∆1 met15∆0 thp1∆::KanMX4* | EUROSCARF |
| FY391 | *MATa ura3∆0 leu2∆0 his3∆1 met15∆0 sac3∆::KanMX4* | EUROSCARF |
| FY395 | *MATa ura3∆0 leu2∆0 his3∆1 met15∆0 cyh2 mex67::KanMX4(DAmP)* | Open Biosystems |
| FY396 | *MATa ura3∆0 leu2∆0 his3∆1 met15∆0 cyh2 mtr2::KanMX4(DAmP)* | Open Biosystems |
| FY399 | *MATa ura3∆0 leu2∆0 his3∆1 met15∆0 sgf11∆::KanMX4* | EUROSCARF |
| FY402 | *MATa ura3∆0 leu2∆0 his3∆1 met15∆0 sgf73∆::KanMX4* | EUROSCARF |
| FY432 (W303R) | *MATa ura3-1 leu2-3,112 trp1-1 his3-11,15 ade2-1 can1-100 RAD5+ RDN1::ADE2* | Won-Ki Huh & Leonard Guarente |
| FY433 (DMY2798) | *MATa ura3-1 leu2-3,112 trp1-1 his3-11,15 ade2-1 can1-100 leu2::mURA3* | Won-Ki Huh & Danesh Moazed |
| FY434 (DMY2804) | *MATa ura3-1 leu2-3,112 trp1-1 his3-11,15 ade2-1 can1-100 RDN1-NTS1::mURA3* | Won-Ki Huh & Danesh Moazed |
| FY435 (DMY2800) | *MATa ura3-1 leu2-3,112 trp1-1 his3-11,15 ade2-1 can1-100 RDN1-NTS2::mURA3* | Won-Ki Huh & Danesh Moazed |
| FY451 | *MATa ura3∆0 leu2∆0 his3∆1 met15∆0 sem1∆::KanMX4* | EUROSCARF |
| FY739 | *MATa ura3∆0 leu2∆0 his3∆1 met15∆0 MEX67-GFP (S65T)::HIS3MX6* | Seong-Tae Kim & Won-Ki Huh |
| FY740 | *MATa ura3∆0 leu2∆0 his3∆1 met15∆0 DBP5-GFP (S65T)::HIS3MX6* | Seong-Tae Kim & Won-Ki Huh |
| SY022 | *MATa ura3∆0 leu2∆0 his3∆1 met15∆0* [pRS316] | This study |
| SY495 | *MATa ura3∆0 leu2∆0 his3∆1 met15∆0 sus1∆::HIS3MX6* | This study |
| SY529 | *MATa ura3∆0 leu2∆0 his3∆1 met15∆0 sir2∆::HIS3MX6* | This study |
| SY532 | *MATa ura3∆0 leu2∆0 his3∆1 met15∆0 sac3∆::KanMX4 sir2∆::HIS3MX6* | This study |
| SY551 | *MATa ura3-1 leu2-3,112 trp1-1 his3-11,15 ade2-1 can1-100 RAD5+ RDN1::ADE2 sir2∆::KanMX4* | [1] |
| SY556 | *MATa ura3∆0 leu2∆0 his3∆1 met15∆0 thp1∆::KanMX4 sir2∆::HIS3MX6* | This study |
| SY557 | *MATa ura3∆0 leu2∆0 his3∆1 met15∆0 ubp8∆::KanMX4 sir2∆::HIS3MX6* | This study |
| SY558 | *MATa ura3∆0 leu2∆0 his3∆1 met15∆0 sgf73∆::KanMX4 sir2∆::HIS3MX6* | This study |
| SY559 | *MATa ura3∆0 leu2∆0 his3∆1 met15∆0 sgf11∆::KanMX4 sir2∆::HIS3MX6* | This study |
| SY575 | *MATa ura3-1 leu2-3,112 trp1-1 his3-11,15 ade2-1 can1-100 leu2::mURA3 sus1∆::KanMX6* | This study |
| SY576 | *MATa ura3-1 leu2-3,112 trp1-1 his3-11,15 ade2-1 can1-100 leu2::mURA3 ubp8∆::KanMX4* | This study |
| SY577 | *MATa ura3-1 leu2-3,112 trp1-1 his3-11,15 ade2-1 can1-100 leu2::mURA3 sgf11∆::KanMX4* | This study |
| SY578 | *MATa ura3-1 leu2-3,112 trp1-1 his3-11,15 ade2-1 can1-100 leu2::mURA3 sgf73∆::KanMX4* | This study |
| SY580 | *MATa ura3-1 leu2-3,112 trp1-1 his3-11,15 ade2-1 can1-100 RDN1-NTS1::mURA3 sus1∆::KanMX6* | This study |
| SY581 | *MATa ura3-1 leu2-3,112 trp1-1 his3-11,15 ade2-1 can1-100 RDN1-NTS1::mURA3 ubp8∆::KanMX4* | This study |
| SY582 | *MATa ura3-1 leu2-3,112 trp1-1 his3-11,15 ade2-1 can1-100 RDN1-NTS1::mURA3 sgf11∆::KanMX4* | This study |
| SY583 | *MATa ura3-1 leu2-3,112 trp1-1 his3-11,15 ade2-1 can1-100 RDN1-NTS1::mURA3 sgf73∆::KanMX4* | This study |
| SY585 | *MATa ura3-1 leu2-3,112 trp1-1 his3-11,15 ade2-1 can1-100 RDN1-NTS2::mURA3 sus1∆::KanMX6* | This study |
| SY586 | *MATa ura3-1 leu2-3,112 trp1-1 his3-11,15 ade2-1 can1-100 RDN1-NTS2::mURA3 ubp8∆::KanMX4* | This study |
| SY587 | *MATa ura3-1 leu2-3,112 trp1-1 his3-11,15 ade2-1 can1-100 RDN1-NTS2::mURA3 sgf11∆::KanMX4* | This study |
| SY588 | *MATa ura3-1 leu2-3,112 trp1-1 his3-11,15 ade2-1 can1-100 RDN1-NTS2::mURA3 sgf73∆::KanMX4* | This study |
| SY652 | *MATa ura3∆0 leu2∆0 his3∆1 met15∆0 sus1∆::HIS3MX6* | This study |
| SY653 | *MATa ura3∆0 leu2∆0 his3∆1 met15∆0 sus1∆::KanMX6* | This study |
| SY654 | *MATa ura3∆0 leu2∆0 his3∆1 met15∆0 sus1∆::KanMX6* | This study |
| SY699 | *MATa ura3∆0 leu2∆0 his3∆1 met15∆0 sus1∆::KanMX6 sir2∆::HIS3MX6* | This study |
| SY783 | *MATa ura3∆0 leu2∆0 his3∆1 met15∆0 sus1∆::HIS3MX6 thp1∆::KanMX4* | This study |
| SY784 | *MATa ura3∆0 leu2∆0 his3∆1 met15∆0 sus1∆::HIS3MX6 sac3∆::KanMX4* | This study |
| SY785 | *MATa ura3∆0 leu2∆0 his3∆1 met15∆0 sus1∆::HIS3MX6 sem1∆::KanMX4* | This study |
| SY786 | *MATa ura3∆0 leu2∆0 his3∆1 met15∆0 sus1∆::HIS3MX6 ubp8∆::KanMX4* | This study |
| SY787 | *MATa ura3∆0 leu2∆0 his3∆1 met15∆0 sus1∆::HIS3MX6 sgf11∆::KanMX4* | This study |
| SY794 | *MATa ura3∆0 leu2∆0 his3∆1 met15∆0 sem1∆::KanMX4* *sir2∆::HIS3MX6* | This study |
| SY797 | *MATa ura3∆0 leu2∆0 his3∆1 met15∆0 sus1∆::HIS3MX6 sgf73∆::KanMX4* | This study |
| SY825 | *MATa ura3-1 leu2-3,112 trp1-1 his3-11,15 ade2-1 can1-100 RAD5+ RDN1::ADE2 sgf11∆::KanMX4* | This study |
| SY826 | *MATa ura3-1 leu2-3,112 trp1-1 his3-11,15 ade2-1 can1-100 RAD5+ RDN1::ADE2 ubp8∆::KanMX4* | This study |
| SY827 | *MATa ura3-1 leu2-3,112 trp1-1 his3-11,15 ade2-1 can1-100 RAD5+ RDN1::ADE2 sgf73Δ::KanMX4* | This study |
| SY831 | *MATa ura3-1 leu2-3,112 trp1-1 his3-11,15 ade2-1 can1-100 RAD5+ RDN1::ADE2 sus1∆::HIS3MX6* | This study |
| SY955 | *MATa ura3∆0 leu2∆0 his3∆1 met15∆0 sus1∆::HIS3MX6* [pRS316-*SUS1*] | This study |
| SY971 | *MATa ura3∆0 leu2∆0 his3∆1 met15∆0 sus1∆::HIS3MX6* [pGP564] | This study |
| SY972 | *MATa ura3∆0 leu2∆0 his3∆1 met15∆0 sus1∆::HIS3MX6* [YGPM2h11] | This study |
| SY973 | *MATa ura3∆0 leu2∆0 his3∆1 met15∆0 sus1∆::HIS3MX6* [YGPM5d22] | This study |
| SY974 | *MATa ura3∆0 leu2∆0 his3∆1 met15∆0 sus1∆::HIS3MX6* [YGPM6e10] | This study |
| SY975 | *MATa ura3∆0 leu2∆0 his3∆1 met15∆0 sus1∆::HIS3MX6* [YGPM25a15] | This study |
| SY976 | *MATa ura3∆0 leu2∆0 his3∆1 met15∆0 sus1∆::HIS3MX6* [YGPM17i23] | This study |
| SY981 | *MATa ura3∆0 leu2∆0 his3∆1 met15∆0 sus1∆::HIS3MX6* [pRS316-*SUS1*] [pGP564] | This study |
| SY982 | *MATa ura3∆0 leu2∆0 his3∆1 met15∆0 sus1∆::HIS3MX6* [pRS316-*SUS1*] [YGPM25d21] | This study |
| SY983 | *MATa ura3∆0 leu2∆0 his3∆1 met15∆0 sus1∆::HIS3MX6* [pRS316-*SUS1*] [YGPM11d19] | This study |
| SY984 | *MATa ura3∆0 leu2∆0 his3∆1 met15∆0 sus1∆::HIS3MX6* [pRS316-*SUS1*] [YGPM11d24] | This study |
| SY985 | *MATa ura3∆0 leu2∆0 his3∆1 met15∆0 sus1∆::HIS3MX6* [pRS316-*SUS1*] [YGPM3j23] | This study |
| SY986 | *MATa ura3∆0 leu2∆0 his3∆1 met15∆0 sus1∆::HIS3MX6* [pRS316-*SUS1*] [YGPM20a24] | This study |
| SY987 | *MATa ura3∆0 leu2∆0 his3∆1 met15∆0 sus1∆::HIS3MX6* [pRS316-*SUS1*] [YGPM20m13] | This study |
| SY988 | *MATa ura3∆0 leu2∆0 his3∆1 met15∆0 sus1∆::HIS3MX6* [pRS316-*SUS1*] [YGPM8e02] | This study |
| SY989 | *MATa ura3∆0 leu2∆0 his3∆1 met15∆0 sus1∆::HIS3MX6* [pRS316-*SUS1*] [YGPM17c06] | This study |
| SY990 | *MATa ura3∆0 leu2∆0 his3∆1 met15∆0 sus1∆::HIS3MX6* [pRS316-*SUS1*] [YGPM11n21] | This study |
| SY991 | *MATa ura3∆0 leu2∆0 his3∆1 met15∆0 sus1∆::HIS3MX6* [pRS316-*SUS1*] [YGPM33c11] | This study |
| SY992 | *MATa ura3∆0 leu2∆0 his3∆1 met15∆0 sus1∆::HIS3MX6* [pRS316-*SUS1*] [YGPM17e06] | This study |
| SY993 | *MATa ura3∆0 leu2∆0 his3∆1 met15∆0 sus1∆::HIS3MX6* [pRS316-*SUS1*] [YGPM28l21] | This study |
| SY994 | *MATa ura3∆0 leu2∆0 his3∆1 met15∆0 sus1∆::HIS3MX6* [pRS316-*SUS1*] [YGPM11j07] | This study |
| SY995 | *MATa ura3∆0 leu2∆0 his3∆1 met15∆0 sus1∆::HIS3MX6* [pRS316-*SUS1*] [YGPM2k17] | This study |
| SY996 | *MATa ura3∆0 leu2∆0 his3∆1 met15∆0 sus1∆::HIS3MX6* [pRS316-*SUS1*] [YGPM13j07] | This study |
| SY997 | *MATa ura3∆0 leu2∆0 his3∆1 met15∆0 sus1∆::HIS3MX6* [pRS316-*SUS1*] [YGPM13p14] | This study |
| SY998 | *MATa ura3∆0 leu2∆0 his3∆1 met15∆0 sus1∆::HIS3MX6* [pRS316-*SUS1*] [YGPM29i02] | This study |
| SY999 | *MATa ura3∆0 leu2∆0 his3∆1 met15∆0 sus1∆::HIS3MX6* [pRS316-*SUS1*] [YGPM18m04] | This study |
| SY1000 | *MATa ura3∆0 leu2∆0 his3∆1 met15∆0 sus1∆::HIS3MX6* [pRS316-*SUS1*] [YGPM29m05] | This study |
| SY1001 | *MATa ura3∆0 leu2∆0 his3∆1 met15∆0 sus1∆::HIS3MX6* [pRS316-*SUS1*] [YGPM15p13] | This study |
| SY1002 | *MATa ura3∆0 leu2∆0 his3∆1 met15∆0 sus1∆::HIS3MX6* [pRS316-*SUS1*] [YGPM29n02] | This study |
| SY1003 | *MATa ura3∆0 leu2∆0 his3∆1 met15∆0 sus1∆::HIS3MX6* [pRS316-*SUS1*] [YGPM19j13] | This study |
| SY1004 | *MATa ura3∆0 leu2∆0 his3∆1 met15∆0 sus1∆::HIS3MX6* [pRS316-*SUS1*] [YGPM31a08] | This study |
| SY1005 | *MATa ura3∆0 leu2∆0 his3∆1 met15∆0 sus1∆::HIS3MX6* [pRS316-*SUS1*] [YGPM28j24] | This study |
| SY1006 | *MATa ura3∆0 leu2∆0 his3∆1 met15∆0 sus1∆::HIS3MX6* [pRS316-*SUS1*] [YGPM16d05] | This study |
| SY1007 | *MATa ura3∆0 leu2∆0 his3∆1 met15∆0 sus1∆::HIS3MX6* [pRS316-*SUS1*] [YGPM2m11] | This study |
| SY1008 | *MATa ura3∆0 leu2∆0 his3∆1 met15∆0 sus1∆::HIS3MX6* [pRS316-*SUS1*] [YGPM3j10] | This study |
| SY1009 | *MATa ura3∆0 leu2∆0 his3∆1 met15∆0 sus1∆::HIS3MX6* [pRS316-*SUS1*] [YGPM21b06] | This study |
| SY1010 | *MATa ura3∆0 leu2∆0 his3∆1 met15∆0 sus1∆::HIS3MX6* [pRS316-*SUS1*] [YGPM25c21] | This study |
| SY1011 | *MATa ura3∆0 leu2∆0 his3∆1 met15∆0 sus1∆::HIS3MX6* [pRS316-*SUS1*] [YGPM5f02] | This study |
| SY1012 | *MATa ura3∆0 leu2∆0 his3∆1 met15∆0 sus1∆::HIS3MX6* [pRS316-*SUS1*] [YGPM10e17] | This study |
| SY1013 | *MATa ura3∆0 leu2∆0 his3∆1 met15∆0 sus1∆::HIS3MX6* [pRS316-*SUS1*] [YGPM27n09] | This study |
| SY1014 | *MATa ura3∆0 leu2∆0 his3∆1 met15∆0 sus1∆::HIS3MX6* [pRS316-*SUS1*] [YGPM17n15] | This study |
| SY1015 | *MATa ura3∆0 leu2∆0 his3∆1 met15∆0 sus1∆::HIS3MX6* [pRS316-*SUS1*] [YGPM8d07] | This study |
| SY1016 | *MATa ura3∆0 leu2∆0 his3∆1 met15∆0 sus1∆::HIS3MX6* [pRS316-*SUS1*] [YGPM14m17] | This study |
| SY1017 | *MATa ura3∆0 leu2∆0 his3∆1 met15∆0 sus1∆::HIS3MX6* [pRS316-*SUS1*] [YGPM14h07] | This study |
| SY1018 | *MATa ura3∆0 leu2∆0 his3∆1 met15∆0 sus1∆::HIS3MX6* [pRS316-*SUS1*] [YGPM11b07] | This study |
| SY1019 | *MATa ura3∆0 leu2∆0 his3∆1 met15∆0 sus1∆::HIS3MX6* [pRS316-*SUS1*] [YGPM32o15] | This study |
| SY1020 | *MATa ura3∆0 leu2∆0 his3∆1 met15∆0 sus1∆::HIS3MX6* [pRS316-*SUS1*] [YGPM9h05] | This study |
| SY1021 | *MATa ura3∆0 leu2∆0 his3∆1 met15∆0 sus1∆::HIS3MX6* [pRS316-*SUS1*] [YGPM26i03] | This study |
| SY1022 | *MATa ura3∆0 leu2∆0 his3∆1 met15∆0 sus1∆::HIS3MX6* [pRS316-*SUS1*] [YGPM31j13] | This study |
| SY1023 | *MATa ura3∆0 leu2∆0 his3∆1 met15∆0 sus1∆::HIS3MX6* [pRS316-*SUS1*] [YGPM6e10] | This study |
| SY1024 | *MATa ura3∆0 leu2∆0 his3∆1 met15∆0 sus1∆::HIS3MX6* [pRS316-*SUS1*] [YGPM2p09] | This study |
| SY1025 | *MATa ura3∆0 leu2∆0 his3∆1 met15∆0 sus1∆::HIS3MX6* [pRS316-*SUS1*] [YGPM20m03] | This study |
| SY1026 | *MATa ura3∆0 leu2∆0 his3∆1 met15∆0 sus1∆::HIS3MX6* [pRS316-*SUS1*] [YGPM5d22] | This study |
| SY1027 | *MATa ura3∆0 leu2∆0 his3∆1 met15∆0 sus1∆::HIS3MX6* [pRS316-*SUS1*] [YGPM2h11] | This study |
| SY1031 | *MATa ura3∆0 leu2∆0 his3∆1 met15∆0 DBP5-GFP (S65T)::KlURA3* | This study |
| SY1032 | *MATa ura3∆0 leu2∆0 his3∆1 met15∆0 sus1∆::HIS3MX6* [pRS316] | This study |
| SY1033 | *MATa ura3∆0 leu2∆0 his3∆1 met15∆0 sus1∆::HIS3MX6* [pRS316-MEX67] | This study |
| SY1034 | *MATa ura3∆0 leu2∆0 his3∆1 met15∆0 sus1∆::HIS3MX6* [pRS316-DBP5] | This study |
| SY1035 | *MATa ura3∆0 leu2∆0 his3∆1 met15∆0 SUS1-HA::KlURA3* | This study |
| SY1036 | *MATa ura3∆0 leu2∆0 his3∆1 met15∆0 MEX67-GFP::HA-KlURA3* | This study |
| SY1037 | *MATa ura3∆0 leu2∆0 his3∆1 met15∆0 DBP5-GFP::HA-KlURA3* | This study |
| SY1038 | *MATa ura3∆0 leu2∆0 his3∆1 met15∆0* [pRS425] | This study |
| SY1039 | *MATa ura3∆0 leu2∆0 his3∆1 met15∆0 sus1∆::HIS3MX6* [pRS425] | This study |
| SY1040 | *MATa ura3∆0 leu2∆0 his3∆1 met15∆0 sus1∆::HIS3MX6* [pRS425-SUS1] | This study |
| SY1041 | *MATa ura3∆0 leu2∆0 his3∆1 met15∆0 sus1∆::HIS3MX6* [pRS425-MEX67] | This study |
| SY1042 | *MATa ura3∆0 leu2∆0 his3∆1 met15∆0 sus1∆::HIS3MX6* [pRS425-DBP5] | This study |
| SY1043 | *MATa ura3∆0 leu2∆0 his3∆1 met15∆0 sus1∆::HIS3MX6 DBP5-GFP (S65T)::KlURA3* | This study |
| SY1045 | *MATa ura3∆0 leu2∆0 his3∆1 met15∆0 sac3∆::KanMX4 DBP5-GFP (S65T)::KlURA3* | This study |
| SY1048 | *MATa ura3∆0 leu2∆0 his3∆1 met15∆0 sus1∆::HIS3MX6 sac3∆::KanMX4 DBP5-GFP (S65T)::KlURA3* | This study |

**Supplementary Table 2.** Plasmids used in this study

| Name | Description | Source |
| --- | --- | --- |
| pFA6a-HIS3MX6 | *pBR322 origin, Amp^R^, HIS3MX6, HIS3 gene from S. kluyveri.* | [2] |
| pFA6a-KanMX6 | *pBR322 origin, Amp^R^, KanMX6, KanMX gene containins the known kanr ORF of the E. coli transposon Tn903.* | [2] |
| pFA6a-GFP-KlURA3 | *pBR322 origin, Amp^R^, URA3, GFP tag with URA3 gene* | [3] |
| pFA6a-HA-KlURA3 | *pBR322 origin, Amp^R^, URA3, Triple HA tag with URA3 gene* | [3] |
| pGP564 | *2μ, Kan^R^, LEU2* | [4] |
| YGPM2h11 | *2μ, Kan^R^, LEU2, [YBR109W-A]&, [ALG1], YSA1, SUS1, CYC8, YBR113W, RAD16, [LYS2]&* | [4] |
| YGPM5d22 | *2μ, Kan^R^, LEU2, [DAP1]&, MEX67, YPL168W, REV3, YPL166W, [SET6], [MLH3]&* | [4] |
| YGPM6e10 | *2μ, Kan^R^, LEU2, [CUE5]&, snR62, WHI2, YOR044W, TOM6, DBP5, STD1, [RAT1]&* | [4] |
| YGPM25a15 | *2μ, Kan^R^, LEU2, [YKL187C]* MTR2 ASH1 SPE1 YKL183C-A [LOT5]** | [4] |
| YGPM17i23 | *2μ, Kan^R^, LEU2, [RSM23]* CWC23 SOH1 SCS3 MET13 MON1 RPS2 YGL123C-A NAB2 GPG1 [PRP43]* | [4] |
| pRS316 | *CEN/ARS, Amp^R^, URA3* | [5] |
| pRS316-SUS1 | *CEN/ARS, Amp^R^, URA3, SUS1 ORF with +- 900 bp of UTR* | This study |
| pRS316-MEX67 | *CEN/ARS, Amp^R^, URA3, MEX67 ORF with +- 900 bp of UTR* | This study |
| pRS316-DBP5 | *CEN/ARS, Amp^R^, URA3, DBP5 ORF with +- 900 bp of UTR* | This study |
| pRS425 | *2μ, Amp^R^, LEU2* | [6] |
| pRS425-SUS1 | *2μ, Amp^R^, LEU2, SUS1-HA with 900 bp upstream and 700 bp downstream of UTR* | This study |
| pRS425-MEX67 | *2μ, Amp^R^, LEU2, MEX67-HA with 900 bp upstream and 700 bp downstream of UTR* | This study |
| pRS425-DBP5 | *2μ, Amp^R^, LEU2, DBP5-HA with 900 bp upstream and 700 bp downstream of UTR* | This study |

**Supplementary Table 3.** Mean lifespans and p values for RLS analysis

| Figure | Strain | Strain description | Mean lifespan | P value compared to matched control strain |
| --- | --- | --- | --- | --- |
| Figure 1A | BY4741 | WT | 27.3 | control |
|  | SY495 | *sus1Δ* | 20.2 | 0.0006, *** |
|  | SY652 | *sus1Δ* | 22.8 | 0.0120, * |
|  | SY653 | *sus1Δ* | 19.0 | 0.0001, *** |
|  | SY654 | *sus1Δ* | 20.8 | 0.0009, *** |
| Figure 2A | BY4741 | WT | 19.2 | control |
|  | SY495 | *sus1Δ* | 13.8 | 0.0131, * |
|  | FY231 | *ubp8Δ* | 31.9 | <0.0001, *** |
|  | FY399 | *sgf11Δ* | 23.8 | 0.3267, ns |
|  | FY402 | *sgf73Δ* | 45.4 | <0.0001, *** |
| Figure 2B | BY4741 | WT | 24.2 | control |
|  | SY495 | *sus1Δ* | 18.9 | 0.0002, *** |
|  | FY390 | *thp1Δ* | 7.2 | <0.0001, *** |
|  | FY391 | *sac3Δ* | 9.3 | <0.0001, *** |
|  | FY451 | *sem1Δ* | 22.5 | 0.0623, ns |
| Figure 3A | BY4741 | WT | 23.7 | control |
|  | SY495 | *sus1Δ* | 17.2 | <0.0001, *** |
|  | FY231 | *ubp8Δ* | 31.6 | 0.0020, ** |
|  | SY786 | *sus1Δ ubp8Δ* | 18.8 | 0.0014, ** |
| Figure 3B | BY4741 | WT | 21.9 | control |
|  | SY495 | *sus1Δ* | 17.0 | 0.0051, ** |
|  | FY399 | *sgf11Δ* | 26.3 | 0.0291, * |
|  | SY787 | *sus1Δ sgf11Δ* | 19.1 | 0.2784,ns |
| Figure 3C | BY4741 | WT | 23.5 | control |
|  | SY495 | *sus1Δ* | 17.1 | <0.0001, *** |
|  | FY402 | *sgf73Δ* | 40.4 | <0.0001, *** |
|  | SY797 | *sus1Δ sgf73Δ* | 23.5 | 0.3870, ns |
| Figure 3D | BY4741 | WT | 21.8 | control |
|  | SY495 | *sus1Δ* | 16.7 | 0.0028, ** |
|  | FY391 | *sac3Δ* | 8.5 | <0.0001, *** |
|  | SY784 | *sus1Δ sac3Δ* | 12.5 | <0.0001, *** |
| Figure 3E | BY4741 | WT | 22.7 | control |
|  | SY495 | *sus1Δ* | 18.7 | 0.0157, * |
|  | FY390 | *thp1Δ* | 8.4 | <0.0001, *** |
|  | SY783 | *sus1Δ thp1Δ* | 20.0 | 0.0567, ns |
| Figure 3F | BY4741 | WT | 24.2 | control |
|  | SY495 | *sus1Δ* | 18.6 | <0.0001, *** |
|  | FY451 | *sem1Δ* | 20.6 | 0.0188, * |
|  | SY785 | *sus1Δ sem1Δ* | 10.5 | <0.0001, *** |
| Figure 4A | BY4741 | WT | 29.7 | <0.0001, *** |
|  | SY529 | *sir2Δ* | 12.4 | control |
|  | SY557 | *sir2Δ ubp8Δ* | 11.9 | 0.2598, ns |
|  | SY559 | *sir2Δ sgf11Δ* | 13.9 | 0.1020, ns |
|  | SY558 | *sir2Δ sgf73Δ* | 13.8 | 0.8582, ns |
| Figure 4B | BY4741 | WT | 29.3 | <0.0001, *** |
|  | SY529 | *sir2Δ* | 8.8 | control |
|  | SY653 | *sus1Δ* | 20.4 | 0.0006, *** |
|  | SY699 | *sir2Δ sus1Δ* | 11.0 | 0.0200, * |
| Figure 4C | BY4741 | WT | 23.7 | <0.0001, *** |
|  | SY529 | *sir2Δ* | 13.2 | control |
|  | SY556 | *sir2Δ thp1Δ* | 5.5 | <0.0001, *** |
|  | SY532 | *sir2Δ sac3Δ* | 5.6 | <0.0001, *** |
| Figure 5D | BY4741 | WT | 23.9 | control |
|  | FY395 | *MEX67-DAmP* | 18.3 | 0.0007, *** |
|  | FY396 | *MTR2-DAmP* | 20.8 | 0.0781, ns |
| Figure 5E | SY971 | *sus1Δ +* pGP564 | 9.5 | control |
|  | SY972 | *sus1Δ +* YGPM2h11 (*SUS1*) | 28.3 | <0.0001, *** |
|  | SY973 | *sus1Δ +* YGPM5d22 (*MEX67*) | 16.1 | <0.0001, *** |
|  | SY974 | *sus1Δ +* YGPM6e10 (*DBP5*) | 24.8 | <0.0001, *** |
| Figure 5G | SY1039 | *sus1Δ +* pRS425 | 18.6 | control |
|  | SY1040 | *sus1Δ +* pRS425-*SUS1* | 25.8 | <0.0001, *** |
|  | SY1041 | *sus1Δ +* pRS425-*MEX67* | 23.7 | 0.0023, ** |
|  | SY1042 | *sus1Δ +* pRS425-*DBP5* | 21.7 | 0.0408, * |
| Figure 5I | SY1051 | *sus1Δ +* YGPM2h11 *+* pRS316 | 17.1 | - |
|  | SY1052 | *sus1Δ +* YGPM5d22 *+* pRS316 | 11.9 | control-1 |
|  | SY1053 | *sus1Δ +* YGPM6e10 *+* pRS316 | 13.4 | control-2 |
|  | SY1054 | *sus1Δ +* YGPM5d22 *+* pRS316-*DBP5* | 14.8 | 0.0391, *  (compared to control-1) |
|  | SY1055 | *sus1Δ +* YGPM6e10 *+* pRS316-*MEX67* | 15.4 | 0.1858, ns  (compared to control-2) |

***, *p* ≤ 0.001; **, *p* ≤ 0.01; *, *p* ≤ 0.05; ns, not significant.
