## Supplementary figures and images for "Sus1 maintains a normal lifespan through regulation of TREX-2 complex-mediated mRNA export"

### Supplementary Figure S1

Suji *et al.*, Supplementary Figure S1

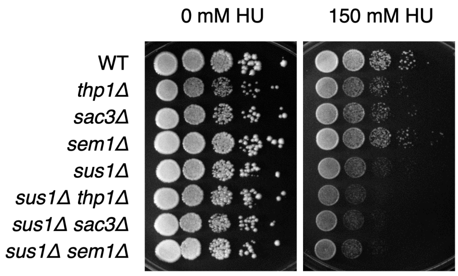

### Supplementary Figure S2

Suji *et al.*, Supplementary Figure S2

A

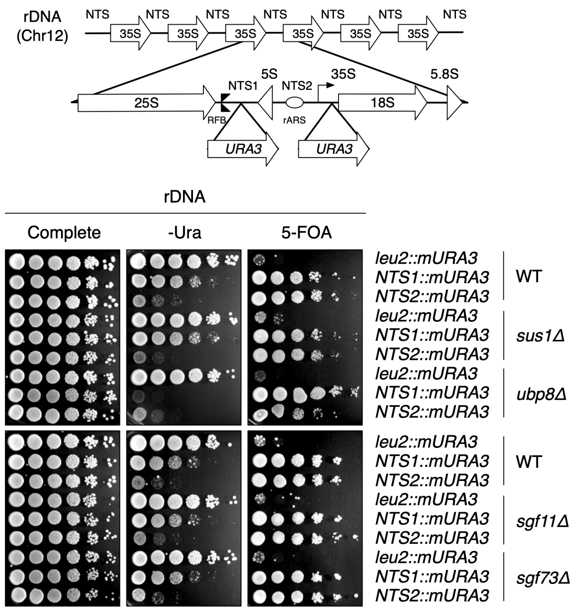


B

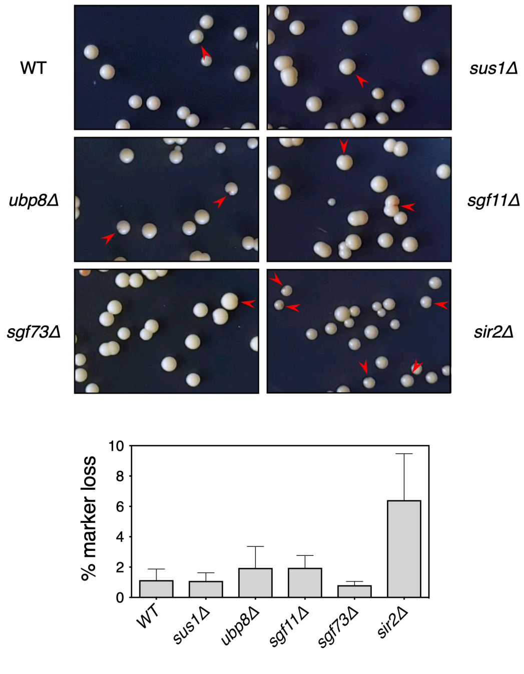

### Supplementary Figure S4

Suji *et al.*, Supplementary Figure S4

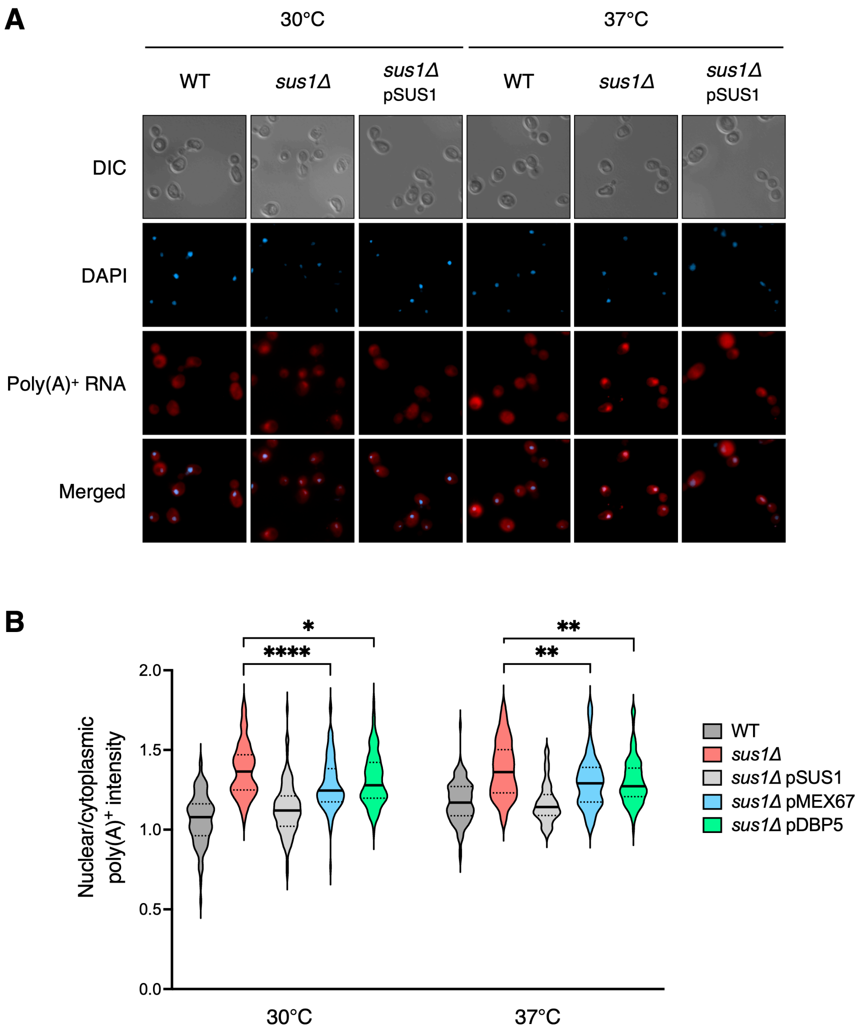
