## Supplementary Figure S3 for "Sus1 maintains a normal lifespan through regulation of TREX-2 complex-mediated mRNA export"

Suji *et al.*, Supplementary Figure S3

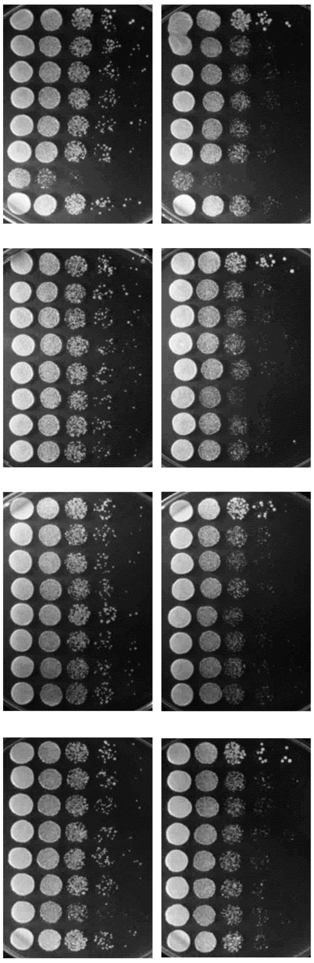


| | Genes in pGP564 |
|---|---|
| pRS316-*SUS1* | - |
| pGP564 | - |
| YGPM25d21 | YAL004W *EFB1 SNR18 VPS8 TFC3* ***NUP60*** |
| YGPM11d19 | ***NUP170*** *ATG8* YBL077W *ILS1* |
| YGPM11d24 | *UGA4 CWC2 NHP2* ***GLE1*** YDL206W *HEM3* |
| YGPM3j23 | YDL121C *YFH1* YDL119C YDL118W *CYK3* ***NUP84*** *IWR1* YDL114W-A YDL114W |
| YGPM20a24 | ***ASM4*** *LUC7* YDL086W YDL086C-A YDL085C-A *NDE2 SUB2 RPS16B* |
| YGPM20m13 | *SLY1 RVB1 HST4* ***NUP42*** YDR193W |
| pRS316-*SUS1* | - |
| pGP564 | - |
| YGPM8e02 | *ARO80 SIP1 CAD1* ***DYN2*** |
| YGPM17c06 | *RTT105* ***NUP157*** *MAM1 GLE2* YER107W-A |
| YGPM11n21 | *SPB4 DEG1 LOC1* ***NIC96*** *YPI1 RPN11* |
| YGPM33c11 | *KEM1* ***NUP49*** *ROK1 SPO74 tK(CUU)G2* |
| YGPM17e06 | *MLC1 ARC1 VPS73 RPL28* YGL102C YGL101W ***SEH1*** |
| YGPM28l21 | *SPC105* ***NUP145*** *NBP35 LIF1* |
| pRS316-*SUS1* | - |
| pGP564 | - |
| YGPM11j07 | YGR117C *RPS23A* ***NUP57*** *COG2 tN(GUU)G MEP1* |
| YGPM2k17 | ***MLP2*** *RPL40A SLN1 ECM37* |
| YGPM13j07 | *HIS5* YIL115W-A ***NUP159*** *POR2 SDP1* |
| YGPM13p14 | *MPM1 DLS1* YJL064W *MRPL8* YJL062W-A *LAS21* ***NUP82*** |
| YGPM29i02 | YJL043W *MHP1* ***NSP1*** |
| YGPM18m04 | ***NUP192*** *tD(GUC)J3 tR(UCU)J2* YJL038C YJL037W *tV(AAC)J* |
| pRS316-*SUS1* | - |
| pGP564 | - |
| YGPM29m05 | *GEF1 URB2* ***NUP85*** |
| YGPM15p13 | YKL070W YKL069W *tV(AAC)K1* YKL068W-A ***NUP100*** *tH(GUG)K YNK1* YKL066W *YET1* |
| YGPM29n02 | *MTD1 RPF2* ***NUP133*** *DAD2 HBS1 MRPL20* |
| YGPM19j13 | *RPL40B* ***MLP1*** *PCC1* |
| YGPM31a08 | ***POM33*** *HIF1 SPA2* YLL020C |
| YGPM28j24 | *GAT3 PPR1 BRE2* YLR016C *MEU1* ***POM34*** |
