## Supplementary Figure S3 (continued) for "Sus1 maintains a normal lifespan through regulation of TREX-2 complex-mediated mRNA export"

Suji *et al.*, Supplementary Figure S3 (continued)

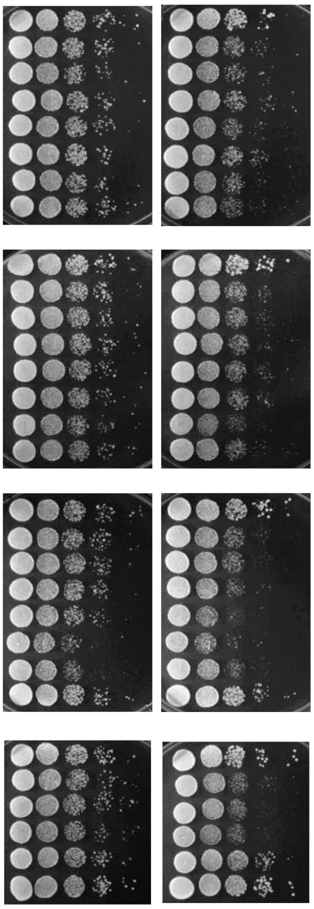


| | Genes in pGP564 |
|---|---|
| pRS316-*SUS1* | - |
| pGP564 | - |
| YGPM16d05 | ***SEC13*** *PNP1* *CLB4* YLR211C |
| YGPM2m11 | *REC102* *CHS5* *JIP3* *MID2* *tD(GUC)L2* *SNR61* *SNR55* *SNR57* *RPS25B* YLR334C *tE(UUC)L* ***NUP2*** |
| YGPM3j10 | ***NUP188*** *CAC2* YML101C-A *CUE4* YML100W-A |
| YGPM21b06 | *AMD1* YML034C-A *SRC1* *RAD52* YML031C-A ***NDC1*** YML030W *USA1* |
| YGPM25c21 | ***POM152*** YMR130W *RRB1* |
| YGPM5f02 | *YIM2* *YIM1* ***NUP53*** YMR153C-A *RIM13* YMR155W *TPP1* *FMP39* *MRPS8* *tM(CAU)M* YMR158C-A YMR158W-B *ATG16* |
| pRS316-*SUS1* | - |
| pGP564 | - |
| YGPM10e17 | *RPS7A* YOR097C ***NUP1*** *KTR1* *CRC1* *RAS1* YOR102W *OST2* |
| YGPM27n09 | *tT(AGU)B* YBR014C *MNN2* YBR016W ***KAP104*** *GAL7* *GAL10* |
| YGPM17n15 | *RMD1* *NTH1* ***YRB1*** *RCR2* YDR003W-A *RAD57* *MAF1* |
| YGPM8d07 | ***SXM1*** YDR396W *NCB2* *UTP5* *HPT1* |
| YGPM14m17 | YER107W-A *FLO8* ***KAP123*** |
| YGPM14h07 | *GUS1* *RTF1* *TAD1* YGL242C ***KAP114*** *DOC1* YGL239C |
| pRS316-*SUS1* | - |
| pGP564 | - |
| YGPM11b07 | YIL067C YIL066W-A *RNR3* *FIS1* YIL064W ***YRB2*** *ARC15* |
| YGPM32o15 | YKL206C ***LOS1*** *EAP1* |
| YGPM9h05 | YLR345W YLR346C ***KAP95*** YLR347W-A *DIC1* YLR349W *ORM2* *NIT3* |
| YGPM26i03 | *REX2* *FRS1* *RPL22A* *BUD28* YLR063W ***PER33*** YLR065C *SPC3* |
| YGPM31j13 | YMR253C YMR254C ***GFD1*** *COX7* *PET111* YMR258C |
| YGPM6e10 | *SNR62* *WHI2* *IRC23* *TOM6* ***DBP5*** *STD1* |
| pRS316-*SUS1* | - |
| pGP564 | - |
| YGPM2p09 | *OSW1* YOR256C ***CDC31*** *HNT3* *RPT4* *GCD1* *RPN8* YOR262W YOR263C *DSE3* |
| YGPM20m03 | YOR111W ***CEX1*** *AZF1* YOR114W *TRS33* |
| YGPM5d22 | ***MEX67*** *MRX4* *REV3* *ATG29* *SET6* |
| YGPM2h11 | *ALG1* *YSA1* ***SUS1*** *CYC8* YBR113W *RAD16* |
