## Supplementary Figure S5 for "Sus1 maintains a normal lifespan through regulation of TREX-2 complex-mediated mRNA export"

Suji *et al.*, Supplementary Figure S5

**A**

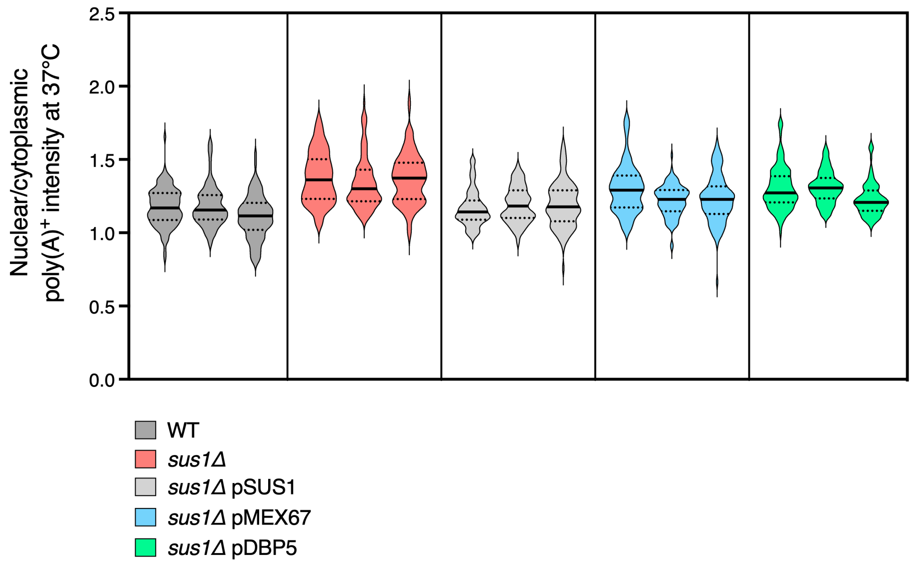


**B**

| Experiments | | Cell numbers | Mean (Nuclear/cytoplasmic poly(A)+ intensity at 37°C) |
|---|---|---|---|
| 1 | WT | 103 | 1.176154 |
| | *sus1Δ* | 112 | 1.376510 |
| | *sus1Δ* pSUS1 | 100 | 1.167284 |
| | *sus1Δ* pMEX67 | 104 | 1.294548 |
| | *sus1Δ* pDBP5 | 103 | 1.304344 |
| 2 | WT | 102 | 1.182704 |
| | *sus1Δ* | 105 | 1.337783 |
| | *sus1Δ* pSUS1 | 105 | 1.188931 |
| | *sus1Δ* pMEX67 | 103 | 1.218946 |
| | *sus1Δ* pDBP5 | 104 | 1.309805 |
| 3 | WT | 103 | 1.107127 |
| | *sus1Δ* | 102 | 1.369916 |
| | *sus1Δ* pSUS1 | 102 | 1.192707 |
| | *sus1Δ* pMEX67 | 110 | 1.231505 |
| | *sus1Δ* pDBP5 | 104 | 1.229178 |
